# Planetary structure and drivers of diazotroph communities reveal key reservoirs of nitrogen-fixation potential

**DOI:** 10.64898/2026.07.27.741065

**Authors:** Lucas J Ustick, Hana Roš, Shahriyar Mahdi Robbani, Jonas Schiller, Anthony Fullam, Anna Mankowski, Chan Yeong Kim, Daniel Podlesny, Kiley Seitz, Thomas S B Schmidt, Samuel Miravet-Verde, Jaime Huerta-Cepas, Shinichi Sunagawa, Michael Kuhn, Peer Bork

## Abstract

Biological nitrogen fixation sustains productivity across Earth’s ecosystems, yet the diversity, distribution and ecological drivers of nitrogen-fixing prokaryotes (diazotrophs) remain incompletely resolved. Here we screened approximately 40 billion genes to generate a catalogue of 21,157 diazotroph genomes. Using this catalogue, we quantified diazotroph diversity and abundance across 9,163 metagenomes spanning diverse environments. Diazotrophy was distributed across the prokaryotic tree of life, but both gene diversity and metagenomic abundance were dominated by heterotrophic lineages. Freshwater and sedimentary ecosystems emerged as major reservoirs of nitrogen-fixation potential beyond the environments that have historically anchored the field. Diazotroph-rich assemblages were associated with nitrogen and phosphorus limitation together with carbohydrate-rich metabolic regimes, revealing distinct nutrient and energy niches for cyanobacterial and heterotrophic diazotrophs. Together, this framework unifies disparate views of diazotroph diversity, abundance and ecology, providing a foundation for improved prediction of biological nitrogen fixation across Earth’s ecosystems.

## Intro

Nitrogen is essential for the synthesis of nucleic acids and proteins, and its availability often limits primary production in terrestrial and aquatic ecosystems (Cui et al. 2025; Du et al. 2020; Moore et al. 2013; Ustick et al. 2021; P. Vitousek and Howarth 1991). Although dinitrogen (N₂) is the most abundant gas in Earth’s atmosphere, it is inert and cannot be assimilated by most living organisms. Biological nitrogen fixation (BNF) accounts for more than 90% of natural nitrogen fixation and is carried out exclusively through nitrogenase enzyme complexes, which convert N_2_ into bioavailable ammonia (Rubio and Ludden 2005; Fowler et al. 2013). Since the early twentieth century, human activities have altered the global nitrogen cycle through industrial nitrogen fixation, fossil fuel combustion, and agricultural intensification, generating reactive nitrogen inputs that rival natural BNF in current estimates (Galloway et al. 2021; Tian et al. 2022).

Despite its importance, the global structure of nitrogen fixing communities remains incompletely resolved. Current views of diazotroph diversity and biogeography derive largely from studies focused on individual environments, specific taxonomic groups, and/or surveys of the marker gene *nifH* (Carpenter and Romans 1991; Karlusich et al. 2021; Xu et al. 2024; Masuda et al. 2024). However, *nifH* has varying copy number per genome, can be difficult to distinguish from homologous sequences and pseudogenes, and does not by itself resolve whether the full genetic machinery required for nitrogen fixation is present (Mise et al. 2021). In addition, prevailing views of diazotroph ecology have been shaped by classic model organisms, especially rhizobia and marine cyanobacteria, and by historical emphasis on open ocean and soil systems (Marcarelli et al. 2022; Turk-Kubo et al. 2023; Capone et al. 1997; Zehr and Capone 2020; Mathesius 2022). These limitations constrain our ability to predict how diazotroph communities will respond to global change. Climate warming is predicted to alter both diazotroph diversity and nitrogen fixation rates (P. Li et al. 2025; Deutsch et al. 2024). At the same time, recent studies suggest that nitrogen fixation may be far more important in coastal margins, freshwater systems, and other previously underrepresented habitats than was assumed in earlier global models (Marcarelli et al. 2022; Fulweiler et al. 2025).

BNF is mediated by three known homologous nitrogenase systems defined by their metal cofactors: the canonical molybdenum-dependent nitrogenase (encoded by *nif* genes) and two alternative systems, the iron-only (*anf*) and vanadium-dependent (*vnf*) nitrogenases (Eady 1996). These systems are typically encoded by conserved catalytic gene operons (*nifHDK, anfHDK, vnfHDK*), which together constitute the minimal genetic machinery required for N_2_ reduction. In many diazotrophs, however, the canonical *nif* operon also includes accessory genes such as *nifE* and *nifN*, which are involved in FeMo-cofactor biosynthesis and nitrogenase maturation. Identification of these conserved operons has enabled systematic detection of diazotrophs from whole-genome data (Bellanger et al. 2024). Recent metagenomic surveys have highlighted that diazotroph communities extend beyond the classical cyanobacterial and rhizobial lineages, with free-living heterotrophic diazotrophs being abundant in much of the open ocean and soils (Delmont et al. 2022; Masuda et al. 2024; Delmont et al. 2018). However, it remains unclear how broadly these patterns extend across Earth’s major ecosystems; which environments serve as principal hotspots of diazotroph diversity and abundance beyond the open ocean and soils; and which ecological drivers consistently structure their distributions within and across biomes.

Here, we present NFixPlanet (nitrogen fixation across the Planet), a global database and cross-biome survey of the nitrogen-fixing microbiome containing 48,151 unique nitrogenase gene sequences, derived from reference genomes (Fullam et al. 2023), MAGs and metagenomic contigs (Schmidt et al. 2024). Using Hidden Markov Models combined with stringent operon-based quality controls, we identified complete molybdenum-, iron-, and vanadium-dependent nitrogenase systems (*nifHDK, anfHDK, vnfHDK*) while explicitly separating them from homologs and pseudogenes. We then mapped an environmentally balanced subset of 9,163 shotgun metagenomes, drawn from an initial pool of 85,604 metagenomes spanning marine, freshwater, terrestrial, anthropogenic and host-associated environments, to this reference framework to quantify diazotroph diversity and global biogeography. Finally, we weighted taxon-resolved diazotroph abundance by published spatial estimates of BNF to identify which lineages dominate nitrogen-fixation potential in environments with high modeled fixation rates. Together, these analyses reveal that diazotrophs are phylogenetically widespread but concentrated within a limited number of major taxa, identify freshwater and sediment systems as hotspots of diazotroph diversity and abundance, link diazotroph-rich communities to nutrient limitation and carbohydrate-rich metabolic regimes, and reveal that heterotrophic lineages dominate nitrogen-fixation potential across biomes.

## Results

### Part 1: NFixPlanet a genome-resolved global reference for diazotroph profiling

We quantified the global genomic potential for nitrogen fixation by screening approximately 40 billion genes derived from high quality reference genomes (proGenomes v3) (Fullam et al. 2023), MAGs and unbinned contigs (SPIRE) (Schmidt et al. 2024) (Figure S1). Nitrogenase genes were identified using Hidden Markov Models (HMMs) combined with stringent genomic context filters designed to distinguish *bona fide* nitrogen-fixation genes from closely related homologs and pseudogenes (Table S1). Only genomes containing the full catalytic gene set (*nifHDK*, *anfHDK*, or *vnfHDK*) within the same genomic neighborhood were considered putative diazotrophs.

Based on these stringent filters we identified 48,151 unique nitrogenase sequences, and 21,157 diazotroph genomes and contigs, including 8,251 reference genomes and 12,906 MAGs and contigs (Figure 1a, 1b, S2). NFixPlanet is the largest collection of diazotroph genomes and genes to date and more than doubles the previously described (Bellanger et al. 2024) whole operon derived diversity of nitrogenase genes introducing 3,834 novel species level *nifD/anfD/vnfD* (97.5% ANI clusters) gene clusters (Figure 1c, Table S2). Leveraging this gene collection, we profiled an environmentally balanced subset of 9,163 shotgun metagenomes drawn from an initial pool of 85,604 metagenomes spanning diverse environments worldwide (Figure 1d, S1). For each metagenome, we quantified diazotroph abundance relative to universal single-copy core genes (Sunagawa et al. 2013) to generate taxon-resolved diazotroph relative abundance profiles. To facilitate reproducibility and future dataset integration, we developed the NFixPlanet Python package and an open web service for identifying nitrogenase genes and profiling metagenomes (nfixplanet.embl.de) (10.5281/zenodo.20644958).

**Figure 1:**
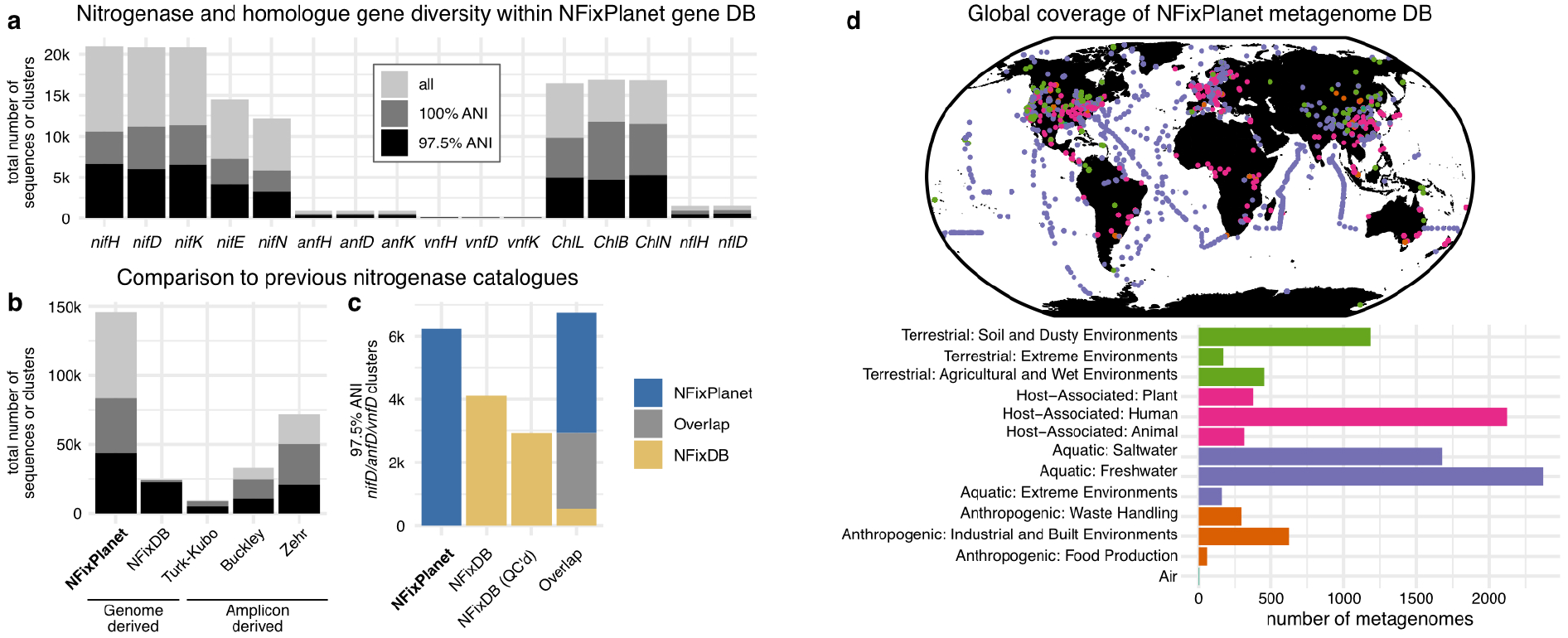
NFixPlanet expands genome-resolved nitrogenase diversity and enables global metagenome profiling. (a) Overview of the NFixPlanet gene DB. Per gene sequence and cluster counts at different average nucleotide identity thresholds for gene clustering. **(b)** Total sequence count per database. NFixPlanet and NFixDB are both derived based on whole-genome sequences while the Turk-Kubo, Buckley, and Zehr databases are all *nifH* and homolog amplicon based. More info about each DB in Table S2. **(c)** NFixPlanet more than doubles the previously described genome derived nitrogenase gene diversity. 97.5% ANI clusters of *nifD/anfD/vnfD* between NFixPlanet and NFixDB. Total number from NFixDB shown followed by the total number of sequences which pass the stringent NFixPlanet quality control steps. Overlap calculated between NFixPlanet and NFixDB sequences which pass QC **(d)** Global distribution of the 9,163 metagenomes selected for mapping systematically subsampled from an initial pool of 85,604 shotgun metagenomes.

### Part 2: Global diversity of nitrogenase genes across the tree of life

Using this dataset, we characterized the richness and phylogenetic distribution of nitrogenase genes across the prokaryotic tree of life. For species-level clustering (97.5% ANI for *nif* genes), we used the catalytic D-subunit genes (*nifD/anfD/vnfD*) rather than *nifH* because *nifD* is single copy while *nifH* can vary in copy number, is more prone to homolog and pseudogene ambiguity (Mise et al. 2021), and, in our benchmarking, recapitulated GTDB species assignments less accurately than *nifD* (Figure S3). Clustering nitrogenase catalytic genes (*nifD/anfD/vnfD*) at 97.5% average nucleotide identity revealed 6,464 unique gene clusters, of which 2,211 (34.2%) were exclusive to cultured genomes, 3,614 (55.9%) were exclusive to MAGs, and 639 (9.9%) occurred in both datasets, highlighting the extent to which culture-independent approaches expand the known diversity of diazotrophs. (Figure S2). Across clusters in the database, the canonical molybdenum-dependent nitrogenase (*nif*) operon dominated. The most common gene arrangements were *nifHDK* (1,825 clusters), *nifHDKEN* (2,871), and *nifHDKE* (918) (Figure S2). These operon structures encode the catalytic core of nitrogenase together with accessory genes involved in FeMo-cofactor biosynthesis and enzyme maturation, and are widely conserved among characterized diazotrophs. In contrast, alternative nitrogenases were comparatively rare: *anf* and *vnf* operons occurred in only 381 and 61 clusters respectively. Because alternative nitrogenases are less efficient than the canonical molybdenum-dependent enzyme, they are typically expressed only under molybdenum and/or vanadium limitation (Mus et al. 2018). Their restricted distribution therefore suggests that the global genomic capacity for nitrogen fixation remains overwhelmingly centered on the canonical molybdenum-dependent system.

Mapping these gene clusters onto the prokaryotic tree of life revealed that diazotrophy is phylogenetically widespread but unevenly distributed. Most nitrogenase-containing genomes and contigs were resolved at broad taxonomic ranks (96% at phylum level), whereas species-level assignments were less complete (64%). However, this difference in annotation depth alone does not explain the strong rank-dependent pattern: species-level assignment was reduced by approximately one-third, whereas the fraction of taxa containing nitrogenase genes declined nearly tenfold, from 29% of phyla (63/217) to 3% of species (3,643/113,104) in GTDB r220 (Figure 2a) (Parks et al. 2022). Thus, diazotrophy spans a broad range of prokaryotic phyla, representing approximately twice the phylum-level breadth previously reported (Bellanger et al. 2024), but remains concentrated within a comparatively small subset of species. Nearly all identified species (99.3%) encoded the canonical *nif* system, and those carrying alternative nitrogenases (*anf* or *vnf*) typically also retained the canonical *nif* machinery (Figure S2). This pattern is consistent with previous evolutionary models proposing that alternative nitrogenases emerged within *nif*-containing lineages (Boyd et al. 2011). Pseudomonadota accounted for the largest share of nitrogenase genes, representing 37% (2,360 genes) of all clusters identified (Figure 2b). Within this phylum, Gammaproteobacteria (1,519 genes) and Alphaproteobacteria (788 genes) dominated. Among the Alphaproteobacteria, most nitrogenase genes were assigned to Rhizobiales (430 genes), an order that includes many well characterized root-nodule symbionts of terrestrial legumes (Poole et al. 2018). Additional gene richness was contributed by Desulfobacterota (873 genes), Bacillota A (504 genes), Bacteroidota (374 genes), and Cyanobacteriota (260 genes) (Figure 2b). The comparatively lower number of clusters observed in Cyanobacteriota suggests that although cyanobacteria are important diazotrophs in many environments their overall richness of nitrogen-fixation genes is much narrower than that of heterotrophic prokaryotes. We compared the predicted life strategies (Podlesny et al. 2026) of genomes encoding nitrogenase genes and found that most were heterotrophic (2,850 genes), whereas autotrophic and mixotrophic lifestyles were less common (391 and 153 genes, respectively) (Figure 2c). Among archaeal phyla, Halobacteriota had the highest number of gene clusters (188 genes) mainly within the methanogenic classes Methanosarcinia (110 genes) and Methanomicrobia (69 genes). This concentration in Halobacteriota is consistent with previous work showing that archaeal diazotrophy is largely restricted to methanogenic anaerobic lineages (Dong et al. 2022). To account for uneven database representation across phyla, we compared gene cluster counts with phylum representation in the combined proGenomes v3 and SPIRE databases using one-sided Fisher’s exact tests (Table S3). All ten phyla differed significantly from the database background, with eight phyla enriched and Bacillota A and Bacteroidota depleted, indicating that the observed patterns cannot be explained solely by sampling bias.

**Figure 2:**
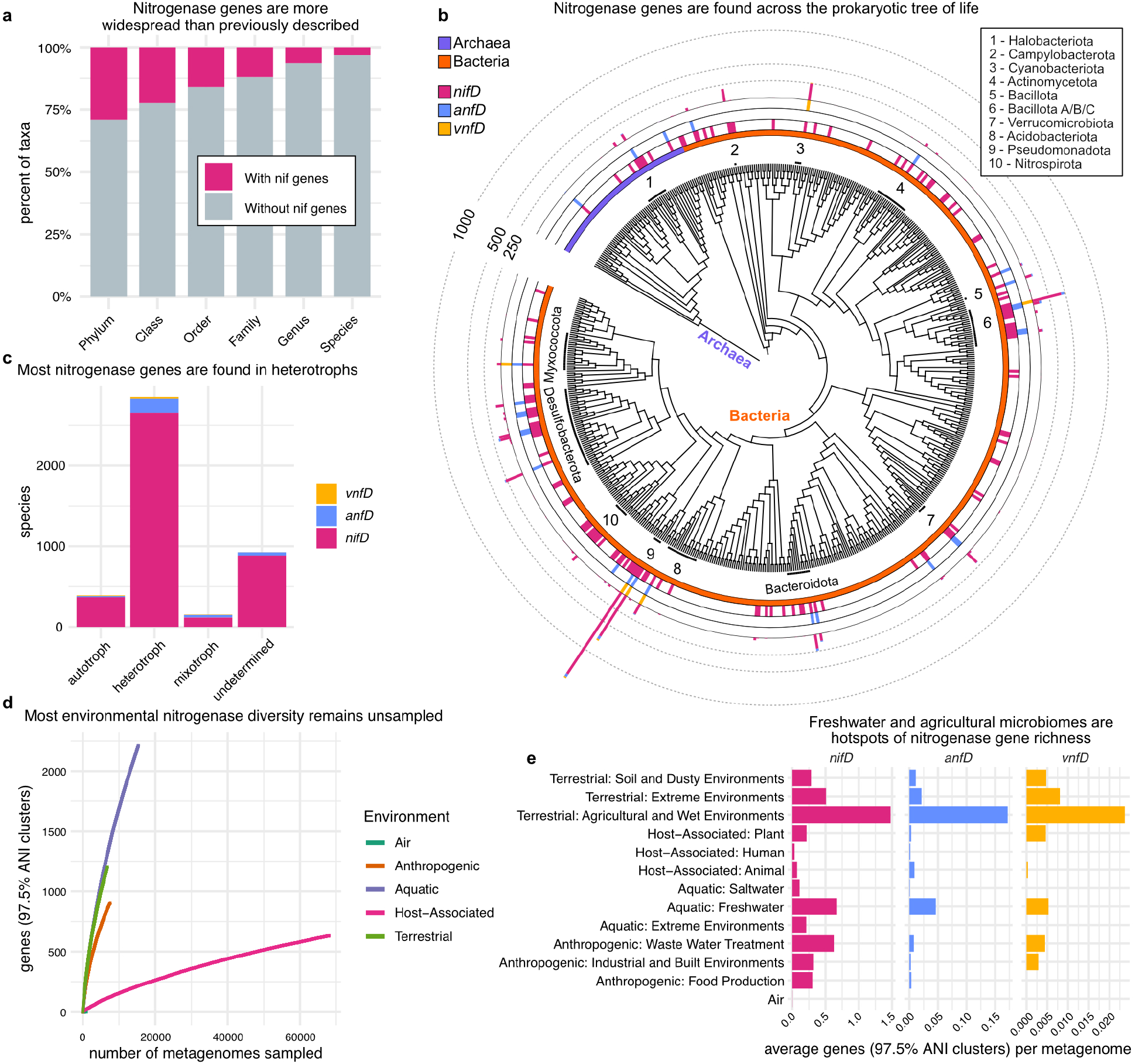
Diazotroph diversity is phylogenetically widespread but concentrated in a limited number of predominantly heterotrophic lineages. **(a)** Percentage of identified taxa at each taxonomic level containing *nifHDK.* (b) GTDB r220 Class-level phylogenetic tree of prokaryotic life showing the distribution and counts of nitrogenase gene clusters. Diazotrophy occurs across a broad span of the prokaryotic tree, but most diversity is concentrated in Pseudomonadota. **(c)** Number of species containing nitrogenase genes grouped based on life strategy. **(d)** Sample-based accumulation analysis of *nifD/anfD/vnfD* gene clusters by environment type. Host-associated communities approached saturation, whereas all other environments remained far from saturation, indicating that substantial diazotroph diversity remains unsampled across all non-host-associated environments. **(e)** Mean nitrogenase gene cluster counts per metagenome across SPIRE-based habitat annotations showing diazotroph diversity hotspots.

Sample-based rarefaction analysis indicated that substantial nitrogenase gene richness remains unsampled across all non-host-associated environments examined (Figure 2d). At a sampling depth of 5,000 metagenomes, aquatic environments showed the greatest increase in accumulated gene-cluster richness (1,055 genes), followed closely by terrestrial (1,007 genes) and anthropogenic (718 genes) environments, whereas host-associated environments showed the smallest increase (97 genes). Agricultural and wetland samples showed the highest mean observed nitrogenase gene cluster richness per metagenomic sample, followed by freshwater and wastewater samples (Figure 2e). In contrast, seawater samples contained substantially fewer observed nitrogenase gene clusters per metagenome than freshwater samples, containing only 15% of the richness per sample compared to freshwater systems.

### Part 3: Global biogeography and environmental drivers of diazotrophs

We quantified diazotroph abundance normalized by total community size (*nifD&K* read coverage / universal single-copy core gene read coverage) by mapping across a balanced subset of 9,163 metagenomes spanning seven ecologically distinct habitat groups (see Methods, Figure 1d, 3a, S4, S5). This number estimates the proportion of the total microbial community which is made up of diazotrophs. The global distribution of diazotroph abundance inferred here broadly matches patterns from available incubation-based rate measurements and biogeochemical models, extending these observations to historically underrepresented environments where comparable data remain scarce. The normalized abundance of diazotrophs was highest in terrestrial hydric systems (median = 4.36%) (including aquifers, sediments, and wetlands) and freshwater environments (median = 1.18%), intermediate in terrestrial soils (median = 0.65%), and lowest in the hot springs (median = 0.07%), anthropogenic environments (median = 0.06%), gut samples (median = 0.05%) and ocean (median = 0.04%) (Figure 3a, 3b). Among aquatic environments, coastal (median = 0.08%) and freshwater systems (median = 1.18%) emerged as global hotspots of diazotroph abundance (Figure 3c). This result aligns with recent syntheses of nitrogen fixation rates derived from incubation experiments, which identify coastal margins as disproportionately important contributors to global nitrogen inputs, suggesting that environments with high diazotroph abundance likely also support elevated fixation activity (Fulweiler et al. 2025). Historically, coastal and riverine environments were expected to exhibit limited diazotrophic activity because of high nutrient inputs from terrestrial sources. Our findings add to a growing body of evidence that these systems support substantial nitrogen fixation (Wen et al. 2017; 2022; Fulweiler et al. 2025). Among human gut samples, preterm infants (median = 3.4%) and individuals with diarrhea (median = 1.8%) had the highest normalized abundance of diazotrophs (Figure S6). Previous work has shown that nitrogen fixation in the human gut occurs at low levels and is unlikely to contribute meaningfully to host nitrogen requirements (Igai et al. 2016). Consistent with this, elevated relative nitrogenase gene abundance was observed only in dysbiotic gut communities, where it is likely driven by depletion of the resident microbiota increasing the relative representation of residual nif-carrying taxa, including opportunistic pathogens such as *Klebsiella,* rather than expansion of nitrogen-fixing populations (McCartney and Hoyles 2023; Mizutani et al. 2021). Diazotroph abundance also co-varied positively with the Shannon diversity of diazotroph populations globally (Pearson’s r = 0.55, p < 2.2e-16; Spearman’s rho = 0.57, p < 2.2e-16; Figure 3d, S7). However, environment-stratified analyses showed that this relationship was weaker within most major biomes, indicating that the global trend is partly driven by biome-level differences in both diazotroph abundance and diversity (Figure S7). Ocean samples occupied the low-abundance, low-diversity end of this gradient, whereas freshwater and terrestrial hydric systems showed higher diazotroph abundance and diversity. This pattern suggests that some environments provide broadly favorable conditions for diazotroph persistence and diversification, while local increases in abundance within a biome may often reflect proliferation of a subset of resident diazotroph taxa rather than expansion of the entire diazotroph assemblage.

**Figure 3:**
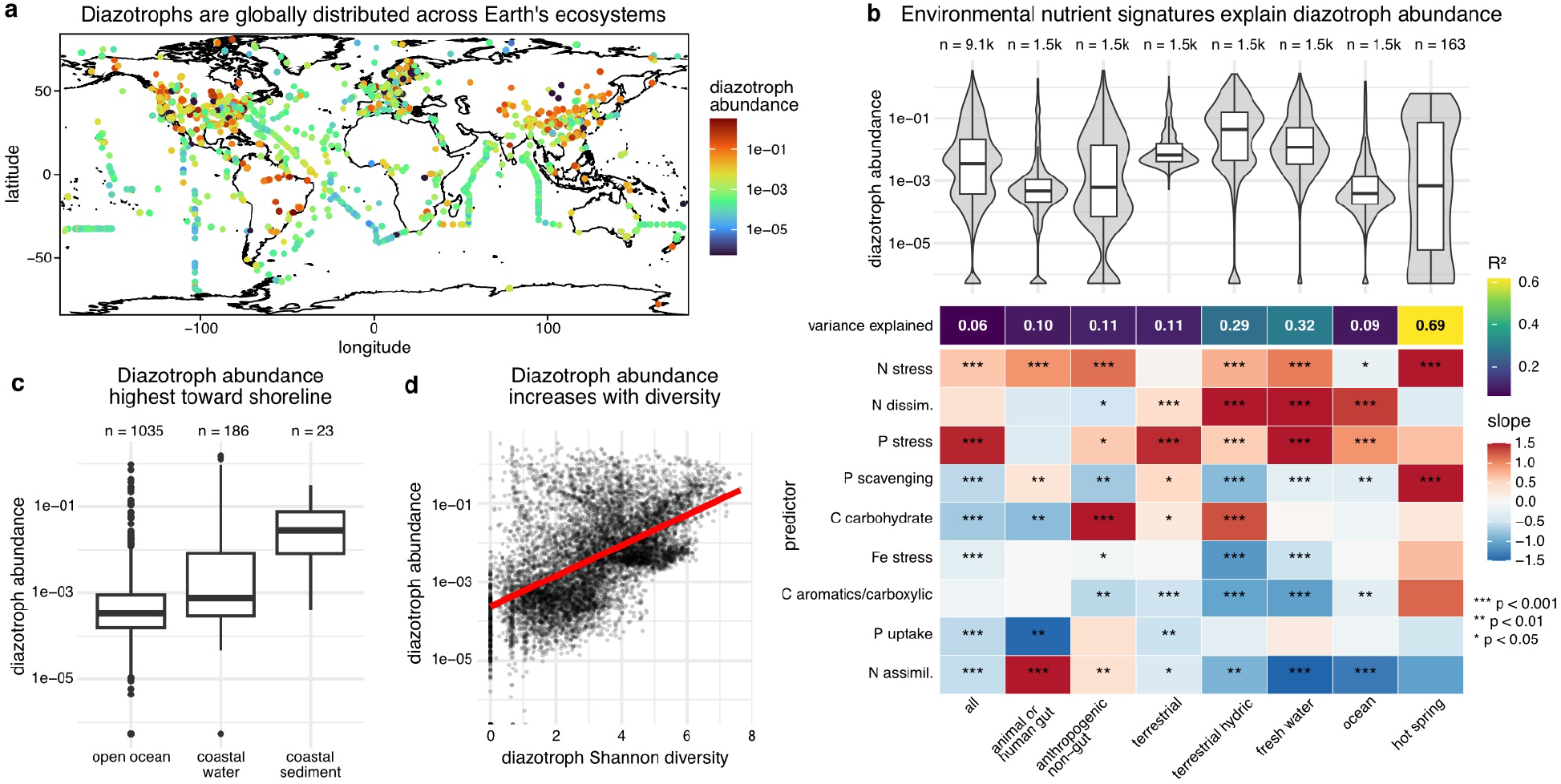
Global cross-environment biogeography of diazotrophs. **(a)** Global distribution of diazotroph normalized abundance across 9,163 metagenomes, calculated as *nifD&K* read coverage normalized by universal single-copy core gene read coverage. This metric approximates the fraction of each microbial community represented by diazotrophs. **(b)** Diazotroph abundance grouped by environment and corresponding linear additive models predicting diazotroph abundance from the whole-community abundance of nutrient-cycling and stress-response genes. Tile color indicates the linear model slope coefficient (effect size); asterisks indicate corresponding Benjamini–Hochberg-adjusted ANOVA significance. Diazotroph abundance is positively associated with signatures of nitrogen stress, phosphorus stress, and carbohydrate metabolism and negatively associated with genes linked to readily available inorganic nitrogen, supporting a model in which diazotrophy is favored both under nitrogen limitation and in carbohydrate-rich environments. **(c)** Increasing diazotroph abundance across the transition from open ocean to coastal waters and coastal sediments. **(d)** Relationship between log-transformed diazotroph abundance and within-diazotroph Shannon diversity across samples (Pearson’s r = 0.55, p < 2.2e-16; Spearman’s rho = 0.57, p < 2.2e-16). Communities with more diazotrophs also tend to contain more diverse diazotroph assemblages.

We quantified the compositional relative abundance of all annotated genes in our metagenomic samples to identify environmental drivers of diazotroph abundance across all biomes. Out of this dataset we selected 115 functional genes related to nutrient cycling and stress responses to model diazotroph abundance, creating linear additive models for all 9,163 metagenomic samples together and each environment split separately (see Methods, Figure 3b, Data S1). Across environments, diazotroph abundance covaried positively with the presence of phosphorus-stress response genes (all lm slope = 1.5, ANOVA p = 1.1*10^-42^), consistent with elevated microbial phosphorus demand in communities where nitrogen fixation supplies additional reduced nitrogen (Figure 3b). Diazotroph-rich communities were characterized by reduced representation of canonical inorganic nitrogen assimilation pathways (all lm slope =-0.56, ANOVA p = 6.9*10^-7^) and increased abundances of nitrogen stress and alternative nitrogen scavenging genes (all lm slope = 0.62, ANOVA p = 4.7*10^-24^), indicating a community-wide shift toward nitrogen acquisition strategies adapted to low inorganic nitrogen availability (Figure 3b). Although iron availability has been proposed as a primary constraint on nitrogen fixation because it is a cofactor of nitrogenase, iron-stress signatures were not among the dominant global predictors of diazotroph abundance across biomes (Berman-Frank et al. 2001; Kustka et al. 2003)(Figure 3b). The two exceptions were in freshwater and coastal/sediment samples, where iron stress was associated with reduced diazotroph representation (freshwater lm slope =-0.53, ANOVA p = 2.4*10^-4^, terrestrial hydric lm slope =-1.12, ANOVA p = 6.4*10^-18^), consistent with strong redox control over benthic iron cycling and iron bioavailability (Burdige and Komada 2020). These associations are consistent with the classical view of diazotrophy as an adaptation to oligotrophic environments where fixed nitrogen is scarce (P. Vitousek and Howarth 1991; Tang et al. 2020; Shen et al. 2024; Moore et al. 2013). Carbon metabolism emerged as an additional axis structuring diazotroph abundance (Figure 3b). Diazotroph abundance was positively associated with genes involved in carbohydrate depolymerization and uptake in anthropogenic, terrestrial, and coastal/sediment samples (mean slope from anthropogenic, terrestrial, and terrestrial hydric models = 1.12, all ANOVA p < 0.05) (Figure 3b). This pattern is consistent with the high energetic demand of nitrogen fixation and with experimental studies showing that bioavailable carbon and energetic supply can stimulate nitrogen fixation, enabling fixation even in nutrient-replete systems (Wang et al. 2021; Dittmann et al. 2025). Conversely, diazotroph abundance declined with increasing representation of pathways associated with aromatic and carboxylic compound degradation in 5 out of 7 of our environmental clusters, highlighting metabolic strategies often linked to oxidative processing of complex organic matter and lower immediate energy yield (Figure 3b). Together, these findings indicate that diazotrophy is favored not only in oligotrophic environments where fixed nitrogen is scarce, but also in energy-rich systems where energetic constraints are alleviated. This energetic coupling provides a plausible mechanistic explanation for the high prevalence of diazotrophs in coastal and sedimentary habitats (Figure 3b, 3c).

### Part 4: Distinct ecological niches of cyanobacterial and heterotrophic diazotrophs

We quantified the taxonomic composition of diazotrophs across the aforementioned 9,163 metagenomes, capturing the planet-wide distribution (Figure 4a). Pseudomonadota were the most abundant diazotroph taxa across all environmental clusters except for hot springs. Cyanobacteriota were found primarily in aquatic environments (Figure 4a, S8). The other most abundant taxa mirrored those with high levels of gene diversity, with Desulfobacterota, Bacillota A, and Bacteroidota being abundant. Halobacteriota were the most abundant archaeal diazotroph taxon found across our samples (Figure 4a). We weighted our in situ taxonomic observations by matching phylum-level diazotroph abundance to spatially resolved global estimates of BNF across terrestrial and open-ocean habitats and to a recent global estimate of inland freshwater BNF (Figure 4b, S9) (Reis Ely et al. 2025; Tang et al. 2019; Fulweiler et al. 2025). This analysis does not directly assign realized fixation rates to taxa, but instead estimates how nitrogen-fixation potential is distributed across environments with different modeled BNF rates. Under this framework, Pseudomonadota represented the largest share of rate-weighted potential (39%) followed by Cyanobacteriota (12%). The role of non-cyanobacterial diazotrophs (NCDs) in aquatic BNF in particular remains unresolved. We found that many Indian Ocean samples were composed almost exclusively of NCDs, including samples from regions where global biogeochemical models predict elevated nitrogen fixation rates (Figures S8, S9). While metagenomic abundance does not directly measure activity, this overlap suggests that heterotrophic diazotrophs may represent an important, and potentially underappreciated, component of marine nitrogen inputs. Together with recent experimental and metagenomic studies, these results support an expanding view of marine diazotrophy in which heterotrophic lineages contribute alongside canonical cyanobacterial taxa (Delmont et al. 2022; Tschitschko et al. 2024).

**Figure 4:**
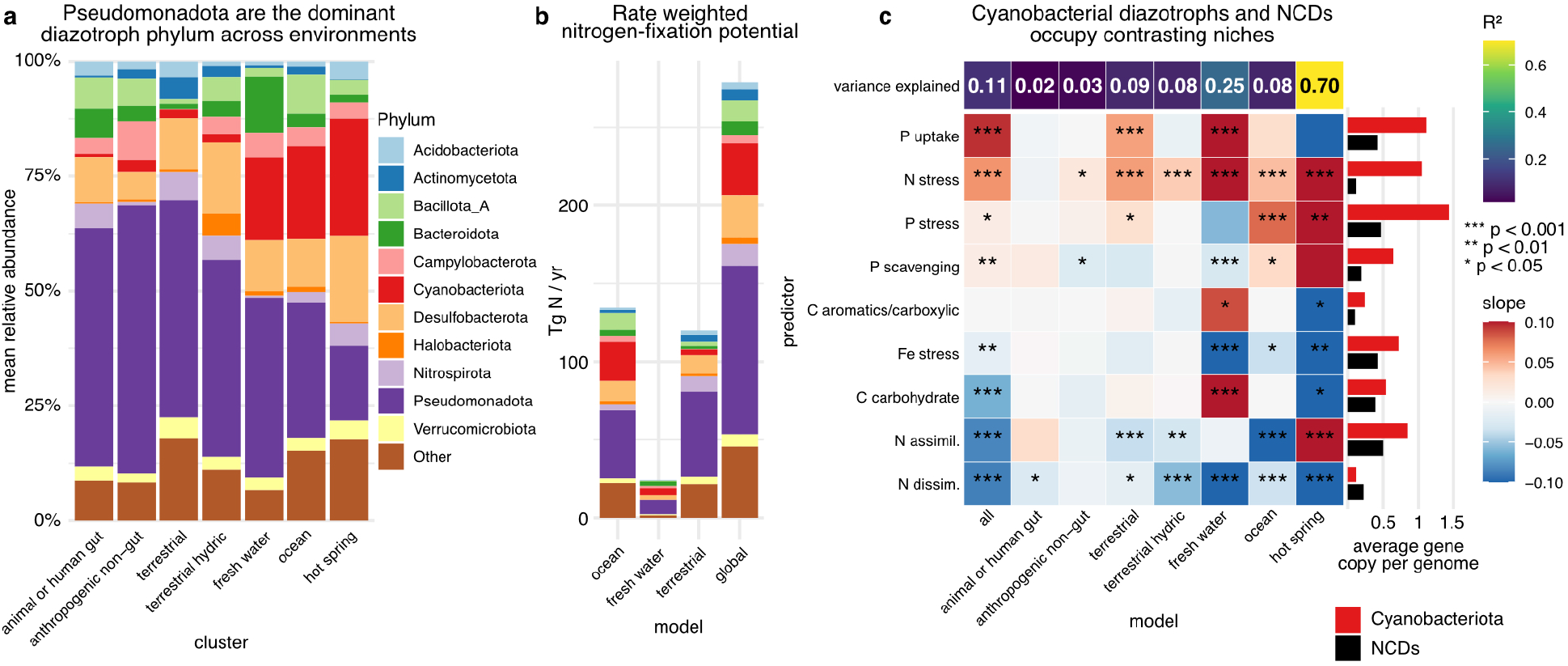
Pseudomonadota dominate global diazotroph abundance, while cyanobacterial and non-cyanobacterial diazotrophs occupy distinct ecological niches. **(a)** Mean phylum-level diazotroph relative abundance across environments. Pseudomonadota dominate in all environments except hot springs, while Cyanobacteriota are mainly found in aquatic systems. **(b)** Rate-weighted nitrogen-fixation potential partitioned by diazotroph phylum, based on published environmental BNF estimates matched with in situ phylum-level diazotroph composition. Pseudomonadota represented the largest share of this rate-weighted potential, followed by Cyanobacteriota. **(c)** Additive linear models relating the relative abundance of cyanobacterial versus non-cyanobacterial diazotrophs (NCDs) to whole-community nutrient-cycling and stress-response gene profiles. Tile color indicates the linear model slope coefficient (effect size); asterisks indicate corresponding Benjamini–Hochberg-adjusted ANOVA significance. Positive coefficients indicate associations with Cyanobacteriota, whereas negative coefficients indicate associations with NCDs. Adjacent bar plots show the mean genomic copy number of genes in each functional category for Cyanobacteriota and NCDs. Cyanobacterial diazotrophs were most strongly associated with functional signatures of nutrient-stressed, oligotrophic conditions, including phosphorus uptake, nitrogen stress, and phosphorus stress. Consistent with these associations, genes linked to these adaptations were more prevalent in cyanobacterial genomes than in NCD genomes. These patterns suggest that cyanobacterial and non-cyanobacterial diazotrophs occupy distinct ecological niches, with cyanobacterial diazotrophy favored under strong nutrient limitation.

We also modeled the relative abundance of cyanobacterial diazotrophs versus NCDs using linear additive models with the aforementioned nutrient cycling genes as predictors (Figure 4c, S8, Data S1). Environmental drivers further differentiate cyanobacterial and NCDs niches. Cyanobacterial diazotrophs were most strongly associated with gene signatures of phosphorus uptake (all lm slope = 0.10, ANOVA p = 1.8*10^-33^) and nitrogen stress (all lm slope = 0.06, ANOVA p = 1.2*10^-73^) and were negatively associated with pathways involved in nitrogen dissimilation (all lm slope =-0.09, ANOVA p = 7.9*10^-45^) and assimilation (all lm slope =-0.08, ANOVA p = 5.1*10^-22^) (Figure 4c). This pattern is consistent with cyanobacterial dominance in oligotrophic environments, where fixed nitrogen is scarce and phosphorus limitation is pronounced (Figure S8). In contrast, model coefficients indicated that NCDs increased relative to cyanobacterial diazotrophs in environments with greater carbohydrate (all lm slope =-0.06, ANOVA p = 2.5*10^-24^) and nutrient availability (Figure 4c). To evaluate whether these environmental associations were reflected in genome content, we quantified the average copy number of nutrient-cycling and stress-response genes across all diazotroph genomes in our dataset. Genes associated with phosphorus uptake, nitrogen stress, and phosphorus stress occurred at an average of more than one copy per Cyanobacteriota genome, whereas these genes were present in less than half of heterotrophic diazotroph genomes on average (Figure 4c). This enrichment of nutrient-stress adaptations in cyanobacterial genomes supports a model in which cyanobacterial diazotrophs are favored under oligotrophic conditions, while NCDs are better able to persist in more energy-rich environments.

## Discussion

A long-standing challenge in nitrogen-fixation research has been to reconcile evidence across levels of organization: from the phylogenetic distribution of nitrogenase genes, to the environmental abundance of diazotrophs, to ecosystem-scale estimates of BNF (Raymond et al. 2004; P. M. Vitousek et al. 2002; 2013). This gap persists in part because the literature has developed unevenly across systems. Marine and plant-associated diazotrophy have been studied intensively, whereas freshwater, sedimentary and coastal microbiomes have been incorporated less consistently into broader syntheses (Zehr and Capone 2020; Fulweiler et al. 2025; Marcarelli et al. 2022; Santi et al. 2013; Mathesius 2022). At the same time, cultivation-independent surveys have shown that nitrogen-fixation potential often resides in taxonomically diverse and frequently heterotrophic lineages that are poorly represented by classical model organisms (Delmont et al. 2018; 2022; Masuda et al. 2024). Our analysis addresses this gap by integrating genome-resolved evidence across major biomes and the prokaryotic tree of life. We find that diazotrophy is broadly distributed phylogenetically, but that its diversity, relative abundance, and cross biome global potential are concentrated disproportionately in heterotrophic groups (Figure 2b, 4a, 4b). We further show that major reservoirs of diazotroph abundance and nitrogen-fixation potential lie in freshwater and sediment microbiomes (Figure 3b, 3c), and thus outside the environments that have historically anchored the field’s conceptual framework. Together, these findings show that major reservoirs of nitrogen-fixing potential span a wider range of environments and taxa than previously assumed, underscoring the need to identify the ecological drivers of diazotroph distributions across ecosystems.

Our results support a view of diazotroph ecology in which nitrogen fixation is shaped by both nutrient limitation and energy supply. Classical theory predicts that diazotrophy should be favoured where fixed nitrogen is scarce, because nitrogenase is energetically costly and is often repressed when reduced nitrogen is readily available (P. Vitousek and Howarth 1991; P. M. Vitousek et al. 2002; Reed et al. 2011; Hoffman et al. 2014). This framework is useful for cyanobacterial diazotrophs: their association with nitrogen and phosphorus stress signatures, alongside the enrichment of nutrient stress genes in Cyanobacteriota diazotroph genomes, is consistent with adaptation to oligotrophic environments (Figure 4c). However, it is insufficient to explain the high diazotroph abundance observed in freshwater, coastal and sedimentary systems, where external nutrient inputs can be substantial (Howarth et al. 1988; Marcarelli et al. 2022; Fulweiler et al. 2025). In such settings, labile carbon and redox structure may act together to favor heterotrophic diazotrophy: organic matter oxidation can supply ATP and reductant, while anoxic or low-oxygen particle and sediment microenvironments protect oxygen-sensitive nitrogenase and maintain the reducing conditions needed for electron transfer to N₂ (Fulweiler 2023; Fulweiler et al. 2007; Dittmann et al. 2025; Wang et al. 2021). The positive association between diazotroph abundance and carbohydrate depolymerization and uptake genes in terrestrial and sediment environments is consistent with this mechanism (Figure 3b). We therefore propose that cyanobacterial and heterotrophic diazotrophs occupy different positions along coupled nutrient, energy and redox gradients: cyanobacteria are favoured where nitrogen scarcity dominates, whereas heterotrophic diazotrophs are favoured where carbon-rich substrates, reducing microenvironments and/or symbiosis reduce the energetic and physiological costs of fixation.

Several caveats should be considered when interpreting these results. First, our analysis quantifies the genetic potential for nitrogen fixation rather than realized fixation rates. Although we implemented stringent filters to distinguish bona fide nitrogenase genes from homologs and pseudogenes, gene presence does not necessarily imply expression or enzymatic activity (Prosser 2015). In particular, our rate-weighted potential analysis (Figure 4b) assumes comparable per-cell fixation rates across taxa and that fixation scales proportionally with relative abundance. This provides a first-order approximation rather than a taxon-resolved flux estimate; physiological differences among diazotrophs, as well as variation in nitrogenase expression, growth state and environmental regulation, may lead to substantial differences in realized fixation rates (Shao et al. 2023). Second, the genome-resolved catalogue itself is shaped by uneven representation of cultivated isolates, MAGs and environments, which may influence inferred patterns of nitrogenase gene diversity. However, Fisher’s exact tests indicated that the phylogenetic composition of the gene catalogue was not explained by database representation alone (Table S3), and these catalogue-level patterns were recapitulated in assembly-free metagenomic profiles: Pseudomonadota contained the largest share of nitrogenase gene clusters and were also the dominant diazotroph phylum across all environmental groups except hot springs (Figure 2d, 4a). Collectively, the agreement between database-level enrichment tests, metagenomic profiles and independent incubation-based syntheses supports the ecological relevance of the trends inferred here (Fulweiler et al. 2025; Shao et al. 2023).

By integrating genome-resolved nitrogenase discovery with cross-biome metagenomic profiling, NFixPlanet expands the view of diazotrophy from a function associated with a limited set of canonical taxa to one distributed across diverse, predominantly heterotrophic lineages. This global perspective identifies freshwater, coastal and sedimentary systems as major reservoirs of nitrogen-fixation potential, and suggests that diazotroph biogeography is shaped not only by nitrogen scarcity but also by carbon availability, nutrient limitation and redox context. Because gene abundance alone cannot resolve realized activity, the next step will be to connect this genomic framework with transcriptomic, proteomic and direct rate measurements across environmental gradients. Beyond sequence-based discovery, approaches leveraging structural similarity may further reveal deeply divergent or currently unrecognized nitrogenase homologs, extending the known functional diversity of diazotrophy. Such integration will be essential for determining when nitrogenase potential is expressed, which diazotrophs contribute most to ecosystem-scale fluxes, and how BNF should be represented in Earth system models under ongoing environmental change.

## Methods

### Initial sequence collection and quality control pipeline

We compiled publicly available Hidden Markov Models (HMMs) for our initial sequence screen. We used the HMMs for *anfHDK*, *ChlLBN*, *nflDH*, *nifHDK*, and *vnfHDK* from (Bellanger et al. 2024). Additionally we included HMMs for *nifEN* from (Aramaki et al. 2020) since they are known homologs of *nifDK*. We ran the aforementioned HMMs against every translated open reading frame (ORF) in the databases proGenomes3 containing approximately 4 billion genes (Fullam et al. 2023) and SPIRE which contains approximately 35 billion genes (Schmidt et al. 2024) using hmmsearch from HMMER v3.4 (http://hmmer.org/)(Eddy 2011). Each ORF was annotated with its top hit, first filtering for the highest bitscore, then the lowest e-value in case of a tie. This helps reduce false positive hits from closely related homologs. For HMMs for *anfHDK*, *ChlLBN*, *nflDH*, *nifHDK*, and *vnfHDK* we quality filtered requiring either an e-value lower than 9.9e-15 or a bitscore larger than 50 following (Bellanger et al. 2024). For *nifEN* we used the suggested bitscore cutoffs of 505.60 and 454.37 respectively (Aramaki et al. 2020).

Because *nifH* and *vnfH* cannot be differentiated by sequences alone, we combined the results from these two HMMs and later re-annotated these two genes based on their proximity to either the *nif* or *vnf* operon.

We further filtered all the HMM results using the following steps. First, we only kept any HMM results which contained the full operon on the same contig. Furthermore we also filtered to make sure all genes were 10 genes up-or downstream of each other in each genome. The sequences retained after this first-pass HMM screen and operon-context filtering are referred to below as the first-pass candidate nitrogenase/homolog sequence set.

### HMM Creation

The public HMMs described above were used as a broad first-pass screen to recover candidate nitrogenase and nitrogenase-homolog sequences from proGenomes3 and SPIRE. We then built custom NFixPlanet HMMs from this expanded first-pass candidate set so that the final models incorporated newly recovered sequence diversity and provided gene-specific gathering thresholds for stringent re-annotation. To reduce redundancy before HMM training, we clustered gene sequences individually at 100% average nucleotide identity using mmseqs easy-cluster from MMseqs2 v14-7e284-gompi-2023a with 80% coverage and bidirectional coverage mode (Steinegger and Söding 2017). The resulting 100% cluster representatives were used as seed sequences to build updated HMMs for each gene with HMMER v3.4. We then calculated bitscore thresholds for each HMM by testing the model against all 100% cluster representatives, treating the seed sequences for the target gene as true positives and all other nitrogenase or homolog sequences as false positives. Following methods described in (Khedkar et al. 2022) we performed 10-fold cross-validation and computed gathering thresholds for each fold as the lowest bitscore at which the highest Matthews Correlation Coefficient was achieved. The final gathering threshold was defined as the median threshold across folds with MCC > 0.8 and was encoded in each HMM profile.

### Final quality control

The final dataset was generated by re-annotating the first-pass candidate nitrogenase/homolog sequence set with the new NFixPlanet HMMs. This first-pass set consisted of all ORFs initially recovered from proGenomes3 and SPIRE using the public HMMs and preliminary operon-context filters, prior to de-replication for HMM training. Candidate ORFs were re-scanned with the NFixPlanet HMMs using hmmsearch with--cut_ga, and the same top-hit assignment and operon-context filters were then reapplied to generate the final curated sequence set.

To validate the classification capability of the pipeline, we evaluated the HMM annotations of the entire final sequence set in terms of clade homogeneity/monophyly via phylogenetic trees (Figure S10-12). All sequences were split into homologous groups (*nifH/nifD/nifK* and corresponding homologs) and de-replicated at 100% average nucleotide identity using MMseqs2 easy-cluster (--cov-mode 0-c 0.8)(Steinegger and Söding 2017). Representative sequences were aligned using MAFFT v7.526 with the FFT-NS-2 algorithm (Katoh and Standley 2013). Alignments were trimmed with trimAl v1.5.1 using a gap threshold of 0.1 (Capella-Gutiérrez et al. 2009). Phylogenetic trees were inferred using FastTree v2.2.0 under default parameters (Price et al. 2010). The trees were visualized in iTOL v7.5.1 (Letunic and Bork 2024), with sequences colored by gene annotations assigned by the NFixPlanet pipeline. Misannotation rates were calculated using tree-traversal functions from ete4 v.4.3.0 (Huerta-Cepas et al. 2016). Each tree was rooted at the midpoint. Each internal node, in a first bottom-up pass, was labeled with the total count of annotations present among all its descendant leaves. In a second top-down pass, each node was assigned a clade label corresponding to the most common annotation in its subtree, switching from the parent label only when a subtree of at least 10 sequences was dominated by a different annotation. This threshold was selected to prevent outliers from redefining clade boundaries. Mismatches were then calculated by comparing each leaf’s clade label against its actual annotation, yielding per-clade counts of incorrectly labeled sequences. Misannotation rates were 9.45 *10^-4^, 4.32*10^-5^, and 1.79*10^-4^ for *nifH*, *nifD*, and *nifK* homologous groups respectively.

### Database comparisons

NFixPlanet was benchmarked against a collection of four publicly available nitrogenase gene data bases (Table S2)(Bellanger et al. 2024; Morando et al. 2025; Gaby and Buckley 2014; Heller et al. 2014). All databases were downloaded, harmonized into comparable FASTA files, and cleaned prior to analysis. We clustered genes in the same manner as previously described using mmseqs easy-cluster from MMseqs2 with 80% coverage and the bidirectional coverage mode. We clustered each database individually at 100% and 97.5% average nucleotide identity. To compare overlap between NFixPlanet and NFixDB (Bellanger et al. 2024) we quality controlled all sequences in NFixDB and removed any sequences which did not pass our filtering steps. We then clustered all *nifD/anfD/vnfD* sequences from NFixPlanet and NFixDB together at 97.5% average nucleotide identity using easy-cluster from MMseqs2 with 80% coverage and the bidirectional coverage mode. We then counted the number of clusters with sequences from only NFixPlanet, only NFixDB, and clusters with sequences from both.

### Universal single copy core gene identification

We identified the 10 mOTUs universal single copy core genes (USCCGs) in both the proGenomes3 and SPIRE databases (Sunagawa et al. 2013). We ran 10 HMMs (COG0012, COG0016, COG0018, COG0172, COG0215, COG0495, COG0525, COG0533, COG0541, COG0552) from fetchMG against every translated ORF in proGenomes3 and SPIRE using hmmsearch from HMMER v3.4 (http://hmmer.org/)(Eddy 2011). We first filtered by the suggested bitscore cutoffs and then again used the top hit method only taking the HMM annotation with the highest bitscore for each ORF. To capture homologs of the 10 SCCGs we also collected any sequences which had a bitscore that fell between 99% and 80% of the gathering threshold and labeled these as false positive homologs. We then clustered the USCCGs and homologs using mmseqs easy-cluster at 50% identity and 80% bidirectional coverage for later usage in the mapping pipeline (Steinegger and Söding 2017). Since we are using these genes to estimate population size regardless of taxonomy we selected this low identity in order to reduce the number of sequences for mapping.

### Taxonomic annotation

All genomes and metagenomic assembled genome bins in the databases proGenomes3 (Fullam et al. 2023) and SPIRE (Schmidt et al. 2024) contained taxonomic assignments based on gtdb-tk v2.4.0 against release r220 (Chaumeil et al. 2022). We removed the annotation for any bins with Domain annotations: “unknown”, “unknown Archaea”, or “unknown Bacteria” which had less than 20% completeness and/or more than 70% contamination as determined by CheckM2 v0.1.3 (Chklovski et al. 2023). These annotations were removed as these are likely false unknowns due to poor assembly/binning. Additionally, any contigs which did not fall into a bin had no taxonomic assignment were annotated against gtdb r220 using CAT_pack pack v6.1beta (von Meijenfeldt et al. 2019).

We assigned life-strategy categories to GTDB r220 species using trait predictions from the metaTraits database (Podlesny et al. 2026). For each species, we extracted predictions for three growth traits: photoautotrophy, photoheterotrophy and chemoheterotrophy. For each trait, species were classified as ‘true’ when the predicted proportion was at least 95%, ‘false’ when it was at most 5%, and ‘mixed’ otherwise. We then defined life strategy from the combination of these trait states. Species were classified as autotrophs when photoautotrophy was ‘true’ and both heterotrophic traits were ‘false’; as heterotrophs when photoautotrophy was ‘false’ and either photoheterotrophy or chemoheterotrophy was ‘true’; and as mixotrophs when photoautotrophy was ‘true’ together with either photoheterotrophy or chemoheterotrophy being ‘true’. All remaining combinations, including species with intermediate (‘mixed’) trait assignments that did not meet these criteria, were classified as undetermined.

We clustered genes at 97.5% ANI as this level most accurately re-create species level taxonomy using *nifH* and *nifD* (Figure S3). We determined this level was the optimum using the following method. We used all sequences with species-level GTDB annotations. We performed clustering (MMseqs2 v. 18.8cc5c; easy-cluster --cov-mode 0,-c 0.8) with percent identity values ranging from 95% to 99% in 0.1% increments, then computed two complementary metrics for each threshold: the mean number of unique species per cluster (USPC), and the mean number of clusters per species (CPS). We used the intersection value of the USPC and CPS curves as the optimum balance between over clustering and underclustering. For both *nifH* and *nifD*, the metrics converged near 97.5% identity, with 93% of clusters having exactly one species annotation (Figure S3).

### Operon completeness filtering

To minimize false classification of incomplete nitrogenase operons caused by contig boundaries when counting operon organizations, we restricted analyses of SPIRE-derived contigs to those with sufficient flanking sequence surrounding the query nitrogenase gene (Figure S2C). For each query, we calculated the maximum absolute neighboring gene offset relative to *nifD*, where the offset represents the number of annotated genes separating a neighboring gene from *nifD*. Only contigs with neighboring annotations extending at least six genes upstream or downstream of *nifD* (maximum absolute offset ≥ 6) were retained. This threshold was selected because *nifE* and *nifN* occurred within six genes of *nifD* in approximately 90% of reference operons, ensuring that their absence on retained contigs most likely reflected genuine operon structure rather than incomplete assembly. Reference genomes from proGenomes3 passed this filtering step.

### Sample selection for mapping

To minimize sampling bias while retaining biologically meaningful environmental variation, we subsampled metagenomes based on previously generated whole-community clustering data (Kim et al. 2026). Briefly, 85,604 metagenomes were clustered using an unsupervised species-composition approach, and all clusters with an overlap coefficient-based phylogenetic distance below 0.34 were grouped into seven ecologically distinct habitat categories (Figure S4): hot spring, ocean, freshwater, terrestrial hydric, terrestrial, anthropogenic non-gut, and animal or human gut. We randomly subsampled 1,500 metagenomes from each category; because only 163 hot spring metagenomes were available, all were retained. This yielded a balanced dataset of 9,163 metagenomes for downstream mapping and analysis.

Marine samples were further subdivided into open ocean, coastal water, and coastal sediment categories (Figure 2C). Samples annotated with both the marine (MICRONT:02020000) and sediment (MICRONT:02070200) microntology terms were classified as coastal sediment (Fullam et al. 2026). For marine water samples, distance to the nearest coastline was calculated using the distancetocoast R package (https://github.com/mdsumner/distancetocoast), and samples within 50 km of the coastline were classified as coastal water or sediment, whereas those located ≥50 km from the coastline were classified as open ocean.

### Mapping pipeline

We compiled a reference database for mapping comprised of all 100% cluster representatives for nitrogenase genes and homologs (*anfHDK*, *ChlLBN*, *nflDH*, *nifHDKEN*, and *vnfHDK*) combined with 50% cluster representatives for all USCCGs and their homologs. We then mapped metagenomes using the following pipeline. Raw metagenomic reads were quality-filtered and trimmed using fastp v0.24.0 (Chen et al. 2018) with sliding-window trimming enabled at both the 5′ and 3′ ends (--cut_front and --cut_tail) using a window size of 4 bp and a mean quality threshold of Q20, and reads shorter than 45 bp were discarded (--length_required 45). Reads were then screened to remove potential host and contaminant sequences using hostile v2.0.0 (Constantinides et al. 2023). The reads were then mapped using CoverM v0.7.0 (Aroney et al. 2025) in contig mode with minimap2 v2.28 (H. Li 2018) using the short-read preset (minimap2-sr). We mapped reads against the reference fasta database. Coverage was calculated using the mean coverage method (--methods mean), and the number of covered bases (--methods covered_bases) was also reported. Both paired-end and unpaired reads were included in mapping where available.

We profiled the mapping results by first summing the coverage gene by gene. For the total abundance of diazotrophs normalized to community size we divided the average coverage of *nifDK* by the average coverage of the universal single copy core genes. For all log-scaled visualizations, abundances were transformed as log(abundance + *m*), where *m* is the minimum non-zero abundance observed in the dataset. This offset was added to retain zero-abundance values during log transformation.

To determine taxonomic relative abundance, we calculated, for each genome, the normalized nitrogenase gene abundance as the sum of the read coverage of *nifDK*, *anfDK*, and *vnfDK* genes divided by the number of these genes detected in that genome (Eq. 1). This produced a genome-level abundance matrix reporting normalized coverage per genome. Genome-level coverage was then summed within each sample according to the taxonomic annotation of each genome across all available taxonomic levels (Eq. 2). To assess the effect of incomplete taxonomic annotation, we quantified the fraction of mapped nitrogenase coverage assigned to references lacking phylum-level taxonomy. Across the seven environment clusters on average, only 0.63–7.56% of mapped nitrogenase coverage was assigned to references without a phylum annotation, indicating that phylum-level diazotroph composition was largely based on taxonomically annotated references.

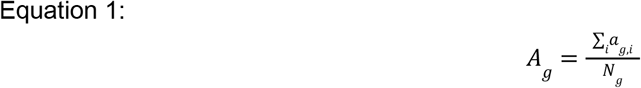

*A_g_*: normalized nitrogenase gene read coverage for genome *g*

*a_g_*_,*i*_: read coverage of gene *i* in genome *g*

*N_g_*: number of nitrogenase genes present in genome *g* (i.e., count of detected genes *i* among *nifDK, anfDK, vnfDK*)

*i*: gene {*nifD,nifK,anfD,anfK,vnfD,vnfK*}

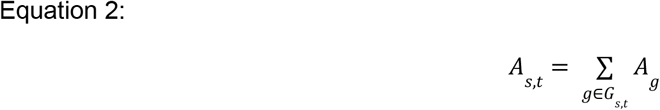

*A_s_*_,*t*_: total gene read coverage for taxon *t* in sample *s*

*A_g_*: normalized nitrogenase gene read coverage for genome *g* (from Eq. 1)

*G_s_*_,*t*_: set of genomes in sample *s* assigned to taxon *t* at a given taxonomic rank

### Linear additive models and gene processing

We created functional profiles for each metagenomic sample by calculating the relative abundance of all ORFs. We included all assemblies with a length greater than 1,000 base pairs and annotated present ORFs using eggNOG-mapper v2.1.3 (Cantalapiedra et al. 2021). We then approximated the abundance of each ORF by multiplying it against the average sequencing coverage of the contig it originated from. ORF coverage was then grouped by KEGG KO (Kanehisa et al. 2004) annotation and made compositional on a per sample basis resulting in the relative abundance of each KEGG KO.

From these profiles, we selected 115 genes related to nutrient cycling and nutrient-limitation stress responses to capture the general nutrient state of each sample (Data S1). To reduce dimensionality and improve interpretability, these 115 KEGG orthologs (KOs) were grouped a priori into biologically coherent functional groups. Specifically, genes were combined into groups describing phosphorus uptake and cycling, phosphorus scavenging, nitrogen dissimilation, nitrogen assimilation, carbohydrate metabolism, aerobic carboxylic/aromatic compound metabolism, iron stress, nitrogen stress, and phosphorus stress. The relative abundances of these genes were then standardized across samples by z-score transformation to place all KOs on a common scale and minimize the influence of differences in baseline abundance among genes. Functional-group-level predictor variables were subsequently calculated as the mean standardized abundance of all KOs assigned to a given functional group.

We modeled diazotroph abundance, and cyanobacterial vs. non-cyanobacterial diazotroph abundance using additive linear regression with the aforementioned gene functional groups as predictors. We employed linear additive models using the lm command from the base R v4.2.1 stats package (https://www.r-project.org/). To evaluate the contribution of each predictor, we performed sequential analysis of variance (ANOVA) on each fitted model using the anova command. Predictors were ordered according to the variance explained by single-predictor linear models (individual R²). p-values from both the linear model coefficient tests and the ANOVA F-tests were adjusted for multiple testing using the Benjamini–Hochberg false discovery rate procedure using the p.adjust command.

To compare the genomic representation of these functional categories between cyanobacterial and non-cyanobacterial diazotrophs, we calculated KO copy number per genome across the selected abundant diazotroph phyla shown in Figure 4a: Acidobacteriota, Actinomycetota, Bacillota_A, Bacteroidota, Campylobacterota, Cyanobacteriota, Desulfobacterota, Halobacteriota, Nitrospirota, Pseudomonadota and Verrucomicrobiota. For each phylum, KO counts were summed across diazotroph genomes, missing phylum–KO combinations were treated as zero, and summed counts were divided by the number of diazotroph genomes assigned to that phylum. KO-level values were then averaged within each functional category. The Cyanobacteriota group corresponds to the category-level values for Cyanobacteriota, whereas the non-cyanobacterial diazotroph group was calculated as the unweighted mean of category-level values across the selected non-Cyanobacteriota phyla.

### Rate-weighted diazotroph potential analysis

To estimate how diazotroph potential is distributed across taxa in environments with different modeled BNF rates, we weighted phylum-level diazotroph relative abundance by published terrestrial, freshwater and ocean BNF estimates. For terrestrial and ocean fixation rates spatially resolved models were leveraged, while for freshwater only global estimates were available. For the two spatially resolved models we summed data into 30 degree bins and multiplied the total fixation within the bin against the average taxonomic composition within the bin (Figure S7). For bins lacking observations, taxonomic composition was inferred from the mean phylum-level total fixation contributions across the observed bins. For the freshwater model we multiplied the global prediction against the global average relative abundance of each phylum. For the terrestrial model, we used the 1° resolution total BNF model from (Reis Ely et al. 2025); for inland freshwater, we used the global predicted BNF from (Fulweiler et al. 2025); and for the ocean, we used the multi-model ensemble mean (n = 13) reported by (Tang et al. 2019).

### Statistical analysis

All statistical analyses were performed using R v4.2.1 (https://www.r-project.org/). To test whether phylum-level differences in nitrogenase gene-cluster richness could be explained by uneven database representation, we compared the distribution of unique *nifD* gene clusters with the distribution of genomes across the combined proGenomes v3 and SPIRE databases for the ten phyla containing the greatest numbers of *nifD* gene clusters (excluding unclassified phyla). One-sided Fisher’s exact tests were performed using fisher.test from the base R stats package, with alternative = “greater” testing enrichment and alternative = “less” testing depletion. All correlations were calculated using the cor.test command from the base R stats package. Sample-based rarefaction analysis was performed separately for each environment using specaccum from the vegan v2.6-2 R package (method = random, 50 permutations) (https://cran.r-project.org/web/packages/vegan/index.html). Shannon diversity of diazotroph taxa was calculated from the genome-level nitrogenase abundance table and normalizing their abundances within each sample. For each sample, genome abundances were converted to within-diazotroph relative abundances, *p_i_* = *a_i_*/Σ*_j_a_j_*, where *a_i_* is the nitrogenase-based abundance of diazotroph genome *i*. Shannon diversity was then calculated as ∑*_i_p_i_log*(*p_i_*) using the diversity function from the vegan package.

### Figure generation

Phylogenetic trees were generated using the Interactive Tree of Life (iTOL) v6 unless otherwise specified (Letunic and Bork 2024). All non-tree figures were generated using ggplot2 in R. Maps were generated using the rnaturalearthdata package (https://cran.r-project.org/web/packages/rnaturalearthdata/index.html) for the topology of land masses.

## Code and Data Availability

Code used to generate this study is available as a python package NFixPlanet v1.0.0 (nfixplanet.embl.de). All data generated in this study is available on Zenodo in the NFixPlanet database release 1.0.0 (10.5281/zenodo.20644958).

## Acknowledgements

Funding and support

This project has received funding from the European Union’s Horizon Europe Research and Innovation Programme under grant agreement no. 101059915 (BIOcean5D). This output reflects only the author’s view, and the European Union cannot be held responsible for any use that may be made of the information contained therein. This work was supported by the EMBL IT Services HPC resources (DOI 10.5281/zenodo.12785829). L.J.U. acknowledges funding from the Peter und Traudl Engelhorn Stiftung zur Förderung der Lebenswissenschaften. S.M.-V. acknowledges funding from the Human Frontier Science Program through the fellowship LT0050/2023-L (https://doi.org/10.52044/HFSP.LT00502023-L.pc.gr.171942).

## Author contributions

L.J.U. and P.B. designed the study. L.J.U. wrote the manuscript with input from all authors. L.J.U., S.M.R., H.R., J.S., A.F., and A.M. processed and quality controlled the data set. L.J.U., S.M.R., and H.R. created and tested the analysis pipeline and computational tools. All authors contributed to the methods development and conceptualization of the analysis. M.K. and P.B. supervised the study.

## Declaration of generative AI and AI-assisted technologies in the writing process

During the preparation of this work, the authors used ChatGPT (OpenAI) in order to assist with language editing, improving clarity, and refining the presentation of the manuscript. After using this tool/service, the authors reviewed and edited the content as needed and take full responsibility for the content of the published article.

## Supplements

**Figure S1:**
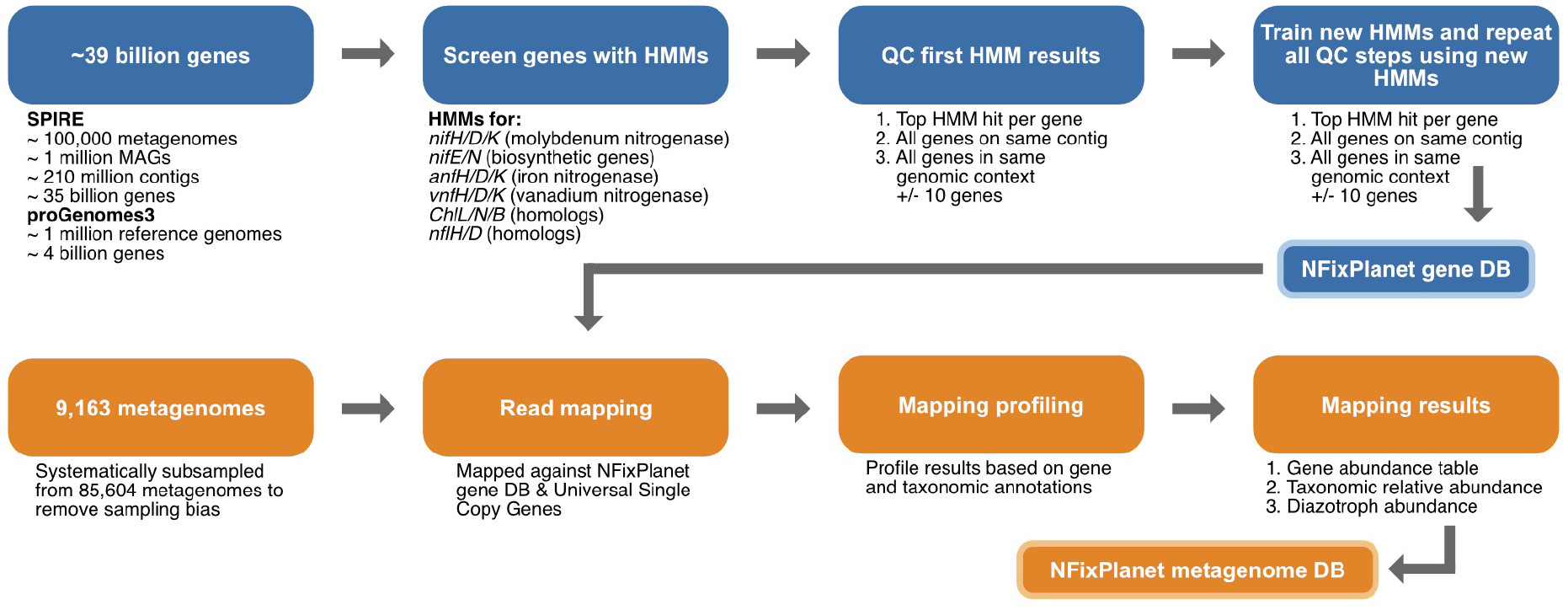
Workflow used to identify genomes and contigs with nitrogenase genes (blue) and the workflow used for profiling metagenomes (orange).

**Figure S2:**
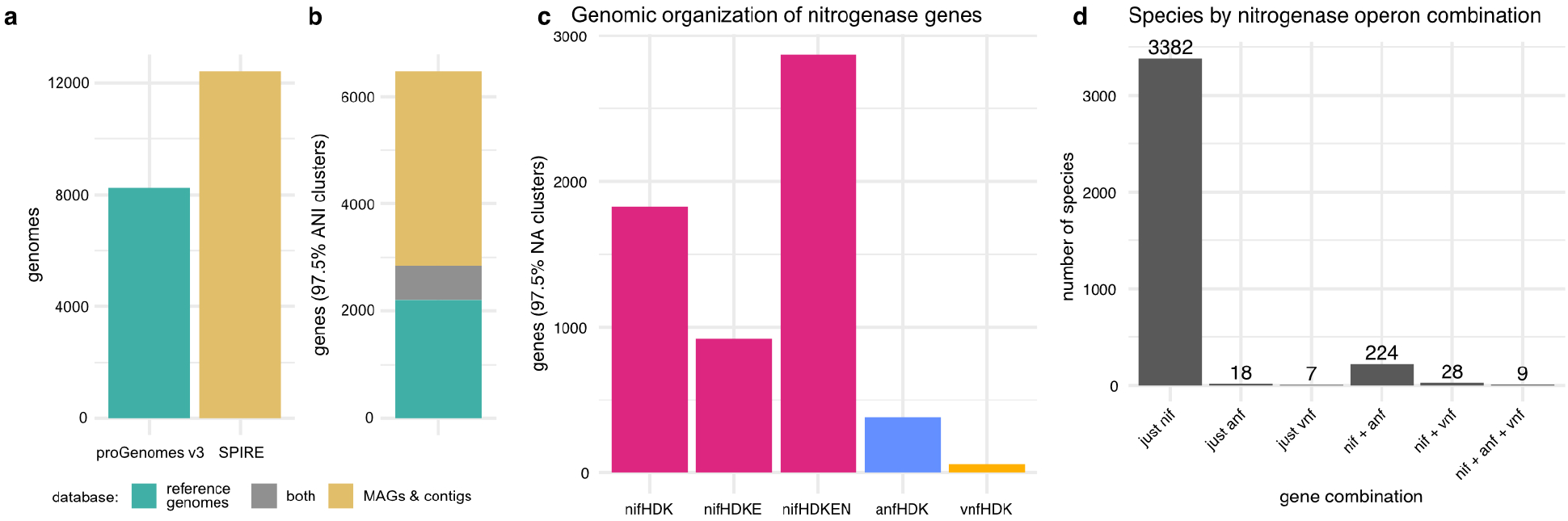
Nitrogenase distributions. **(a)** Total number of diazotroph genomes identified in this study, separated into cultured reference genomes, and metagenome-assembled genomes (MAGs) and contigs. **(b)** Number of *nifD/anfD/vnfD* gene clusters at 97.5% average nucleotide identity, grouped by whether they were found in reference genomes, metagenomic assemblies, or both. **(c)** Counts of distinct genomic arrangements of nitrogenase genes. Canonical Mo-dependent *nif* operons dominate, whereas alternative *anf* and *vnf* operons are comparatively rare. **(d)** Number of species grouped based on which nitrogenase operons are present in the genome.

**Figure S3:**
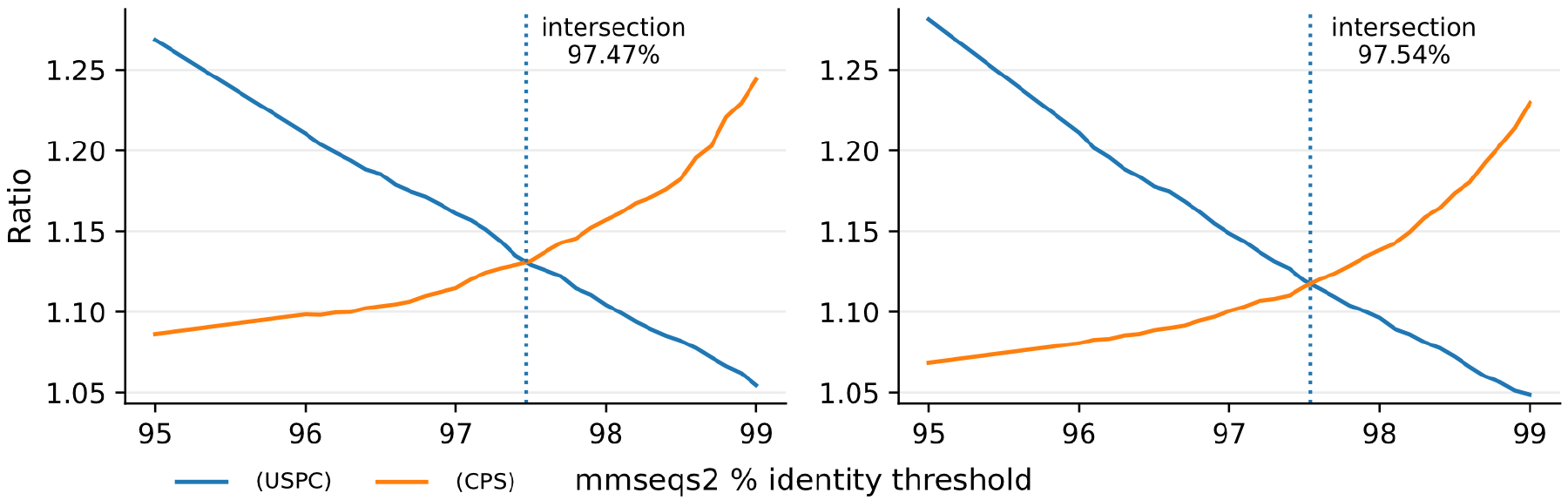
Comparison of clustering performance for nifH and nifD across ANI thresholds. The intersection of unique species per cluster (USPC) and clusters per species (CPS) occurred near 97.5% ANI for both genes, with *nifD* showing better agreement with GTDB species assignments.

**Figure S4:**
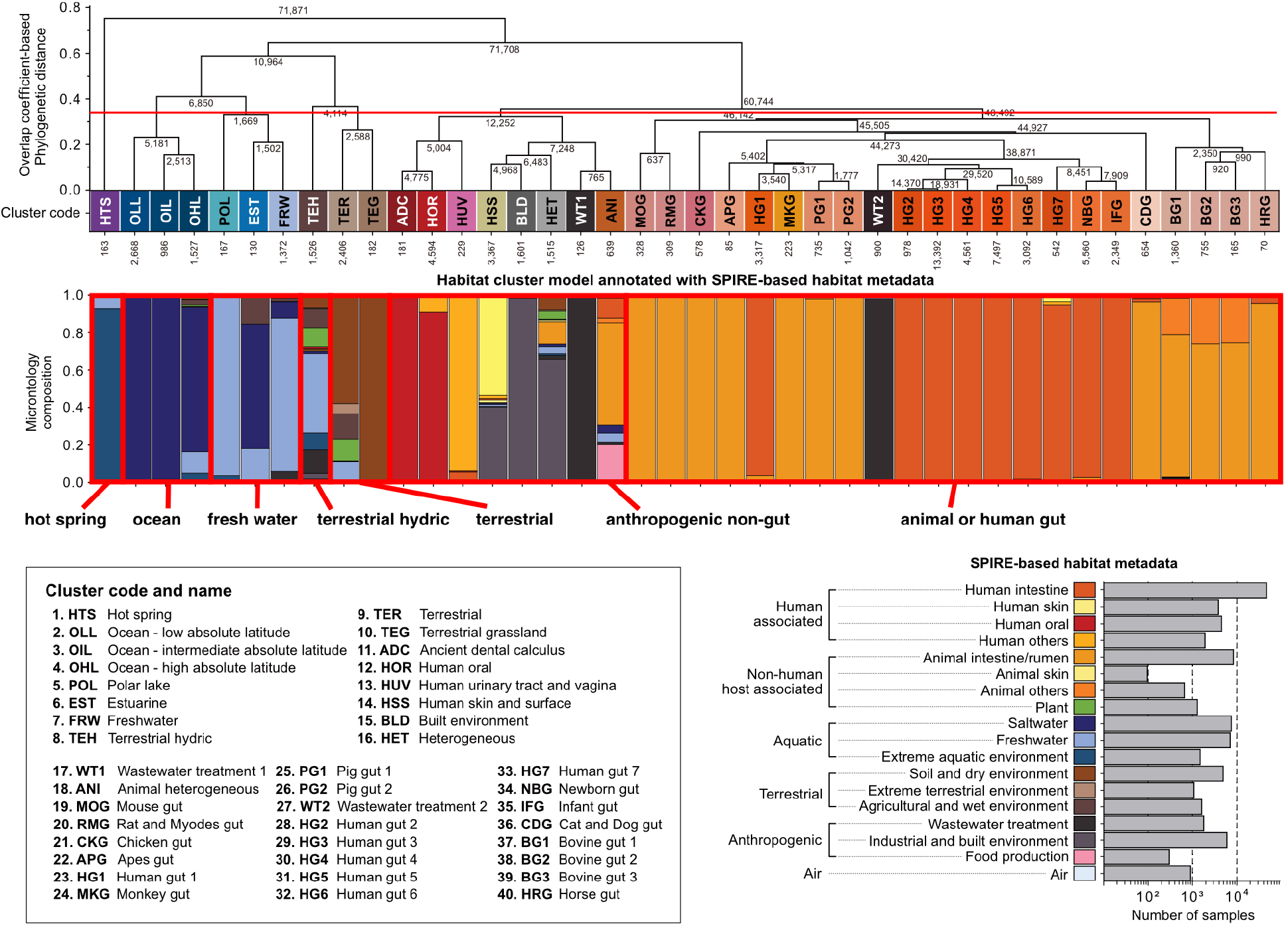
Whole community based clusters of metagenomic samples. Samples were clustered based on whole community composition previously in (Kim et al. 2026). We leveraged this clustering and further grouped the 40 clusters into 7 based on the overlap coefficient-based phylogenetic distance. Within these 7 new clusters we subsampled the metagenomes to have more even coverage across varied environments.

**Figure S5:**
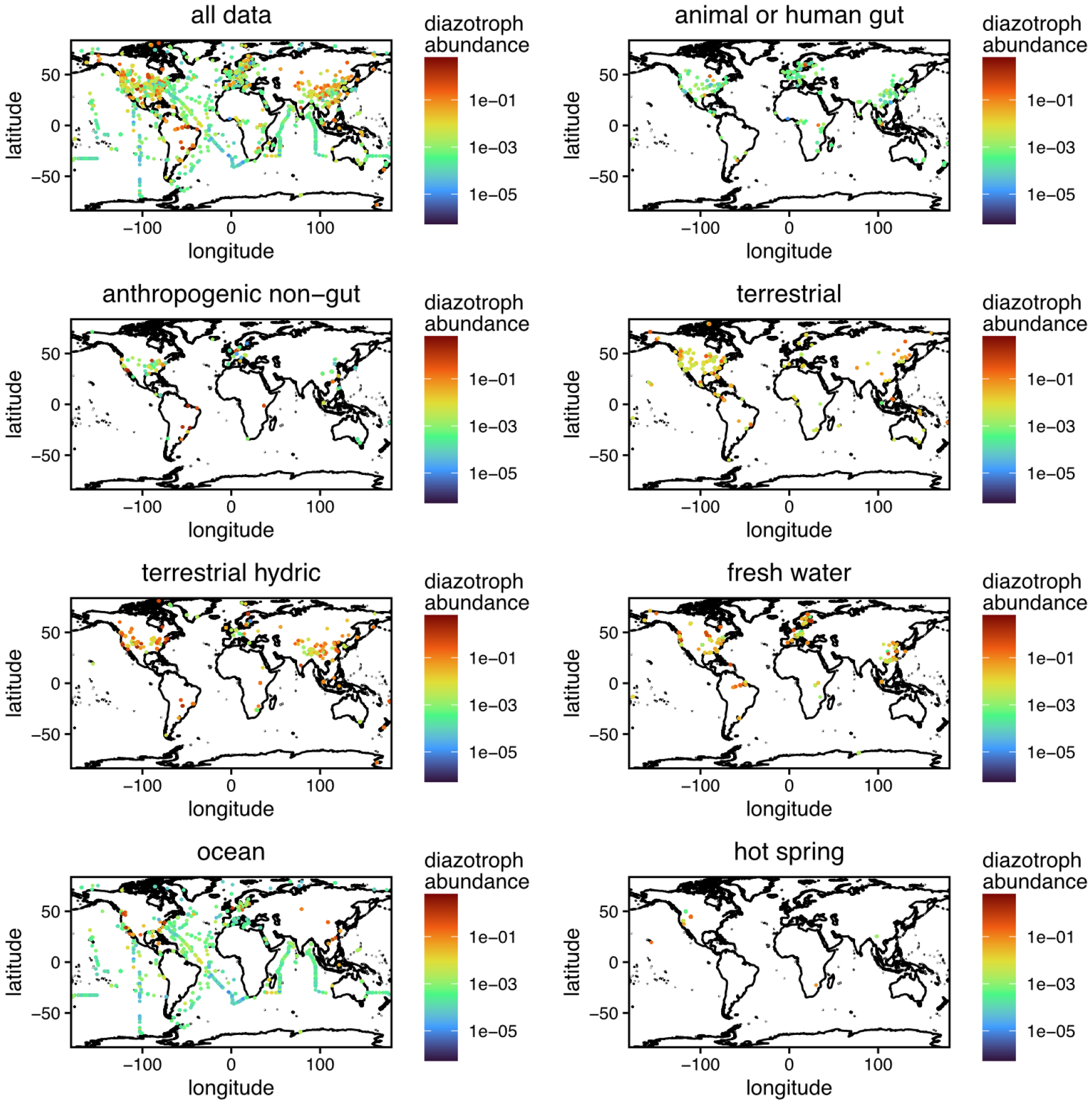
Global distribution of diazotroph abundance across 9,163 metagenomes, calculated as *nifD&K* read coverage normalized by universal single-copy core gene read coverage split by environment. This metric approximates the fraction of each microbial community represented by diazotrophs.

**Figure S6:**
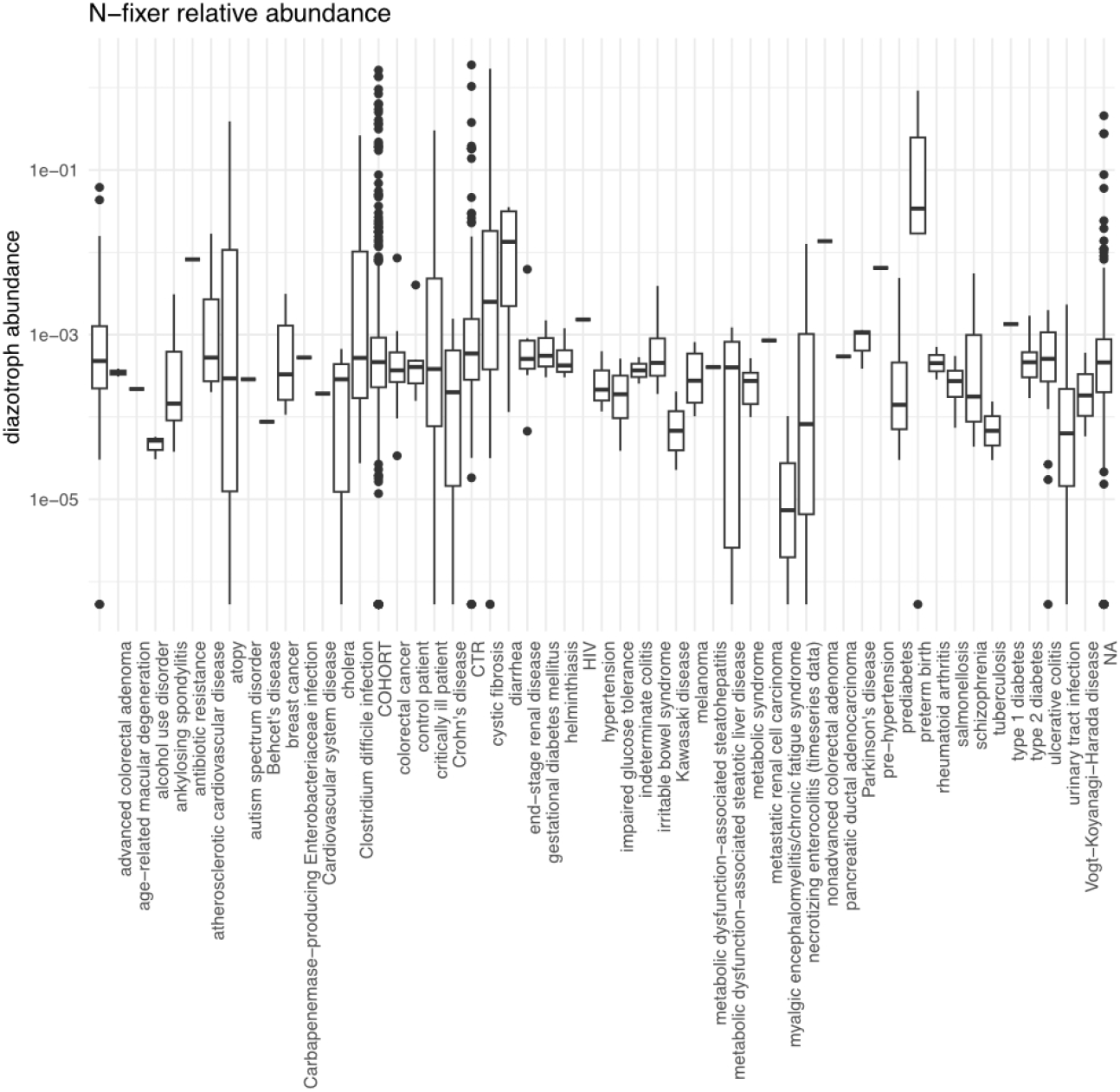
**Relative diazotroph abundance (*nifD&K* read coverage normalized by universal single-copy core gene read coverage) in human gut samples split based on disease status.**

**Figure S7:**
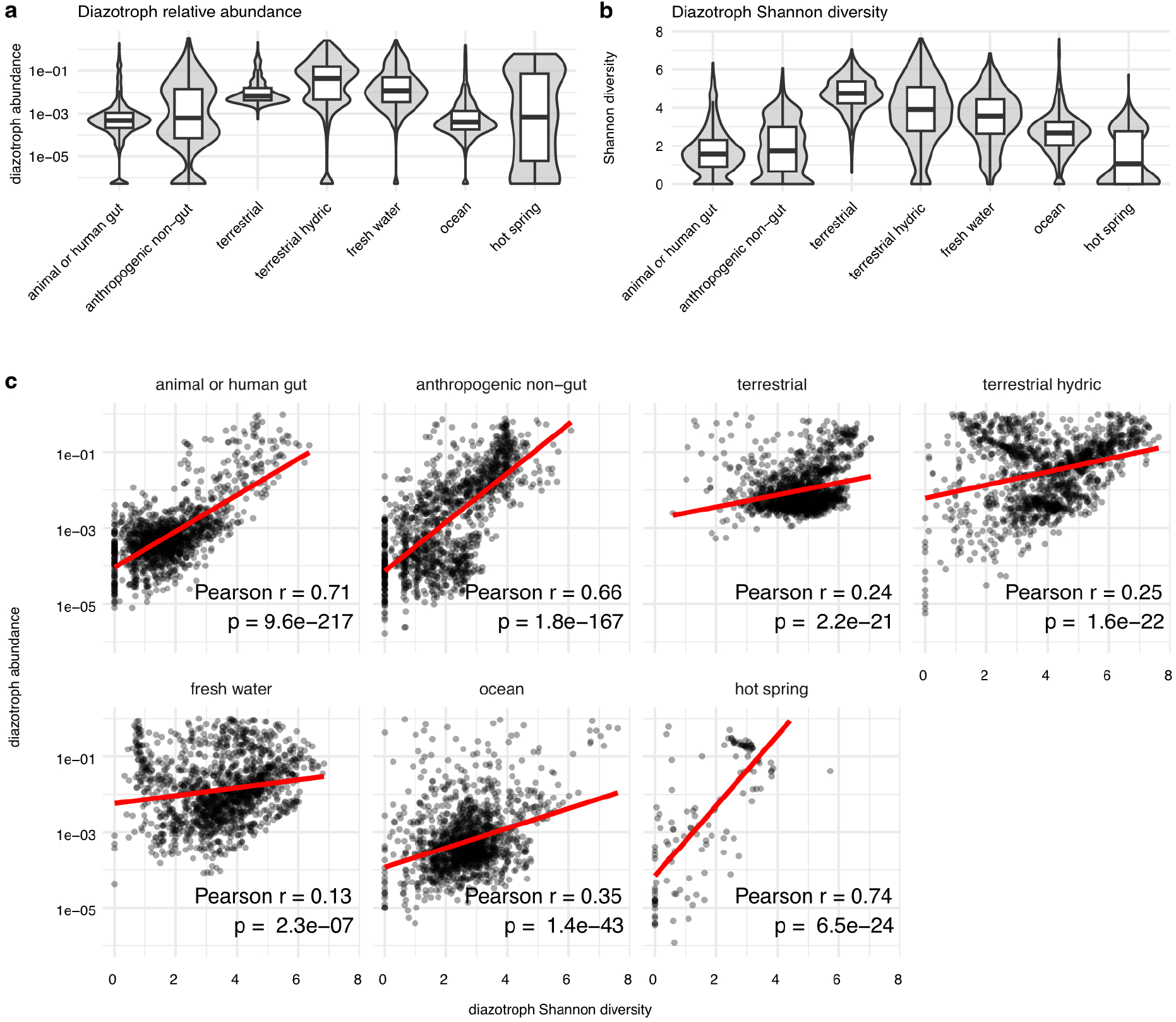
(a) Diazotroph abundance and (b) Shannon diversity of diazotrophs grouped by environment cluster. (c) Relationship between diazotroph abundance and Shannon diversity within each environmental cluster. Diazotroph abundance and diversity remain positively associated within environments, although more weakly than across environments.

**Figure S8:**
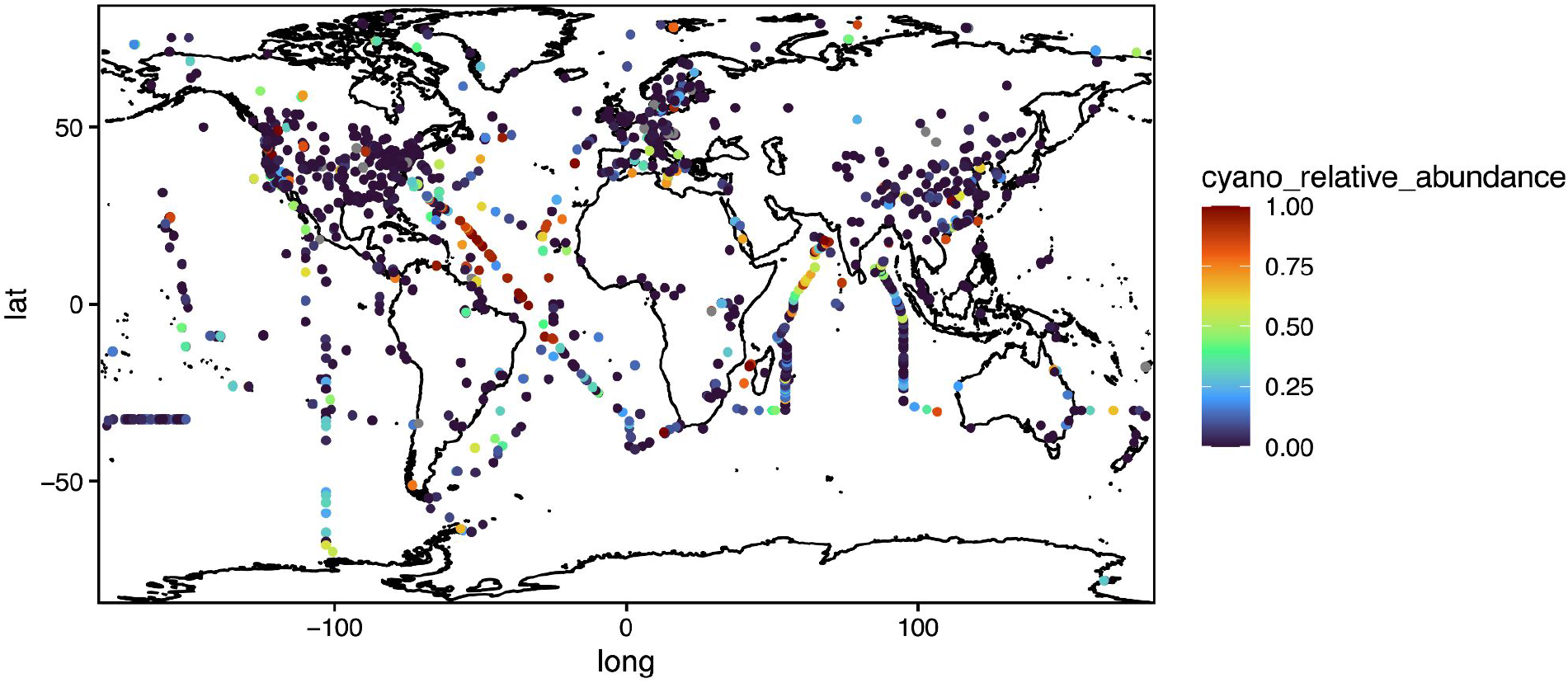
Global distribution of the relative abundance of cyanobacterial versus non-cyanobacterial diazotrophs.

**Figure S9:**
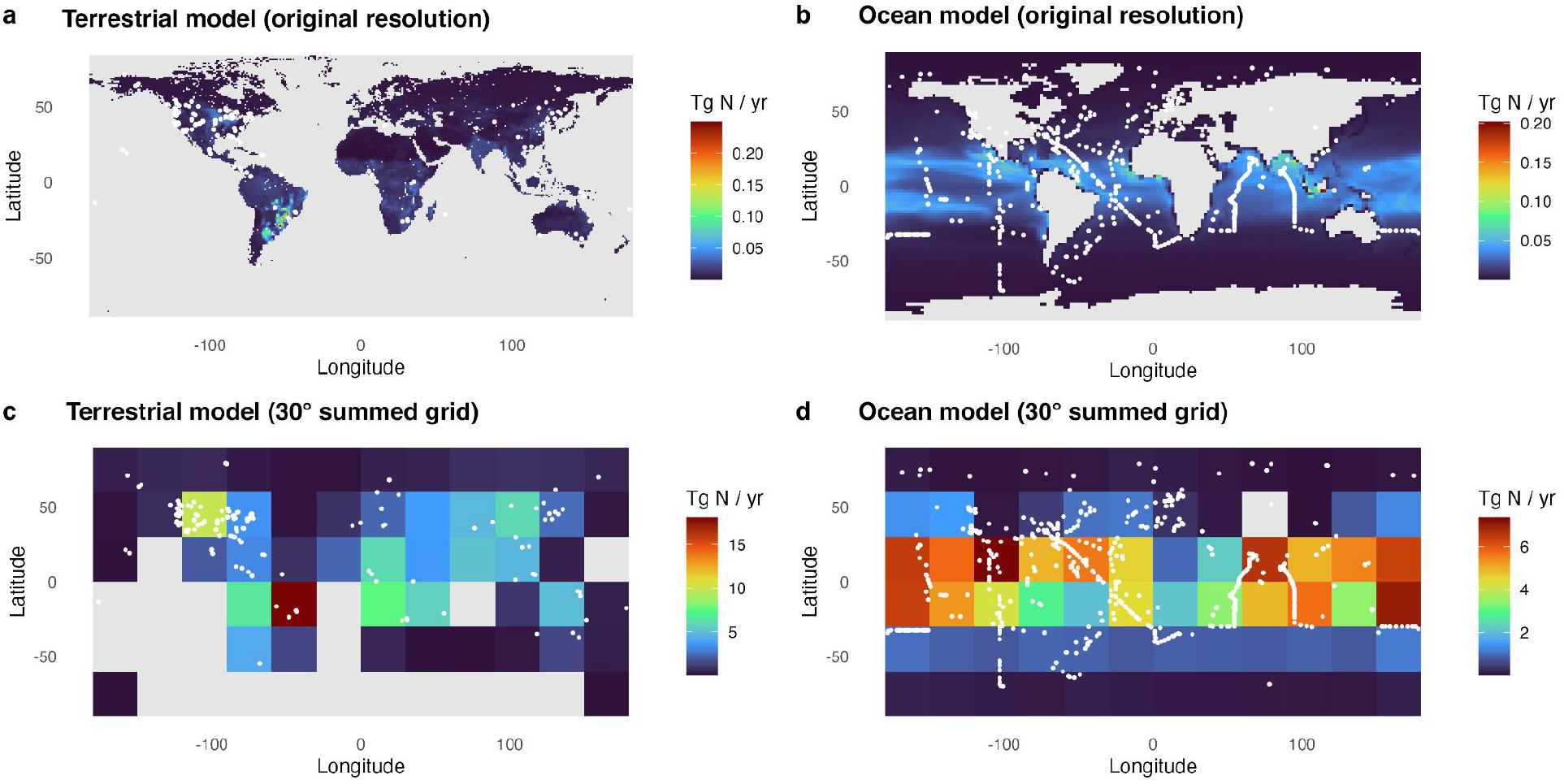
Global nitrogen fixation rate models. **(a)** Original terrestrial model data from (Reis Ely et al. 2025) with metagenomic observations overlaid. **(b)** Original ocean model with metagenomic observations overlaid. Ocean model data is the mean values across 13 global model estimates processed together from (Tang et al. 2019). **(c,d)** Summed and gridded data used for the matchup between metagenomic observations and model data to predict taxon attributed nitrogen fixation.

**Figure S10:**
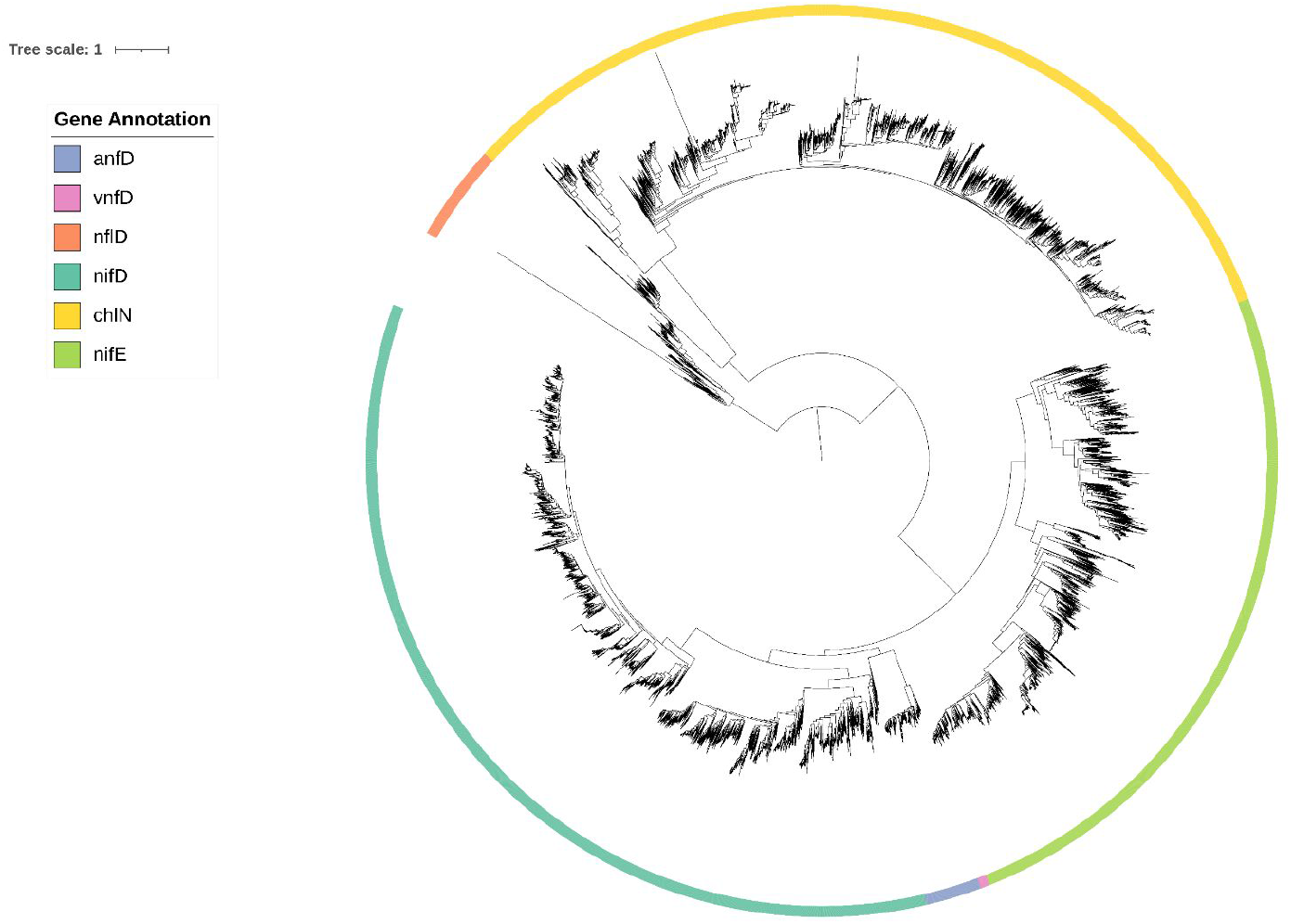
Gene tree of *nifD* and *nifD* homologs. The outer ring indicates final HMM annotations. Tree topology was used to estimate annotation consistency, with one sequence assigned to a non-homogeneous clade (estimated error rate = 4.3 * 10⁻⁵).

**Figure S11:**
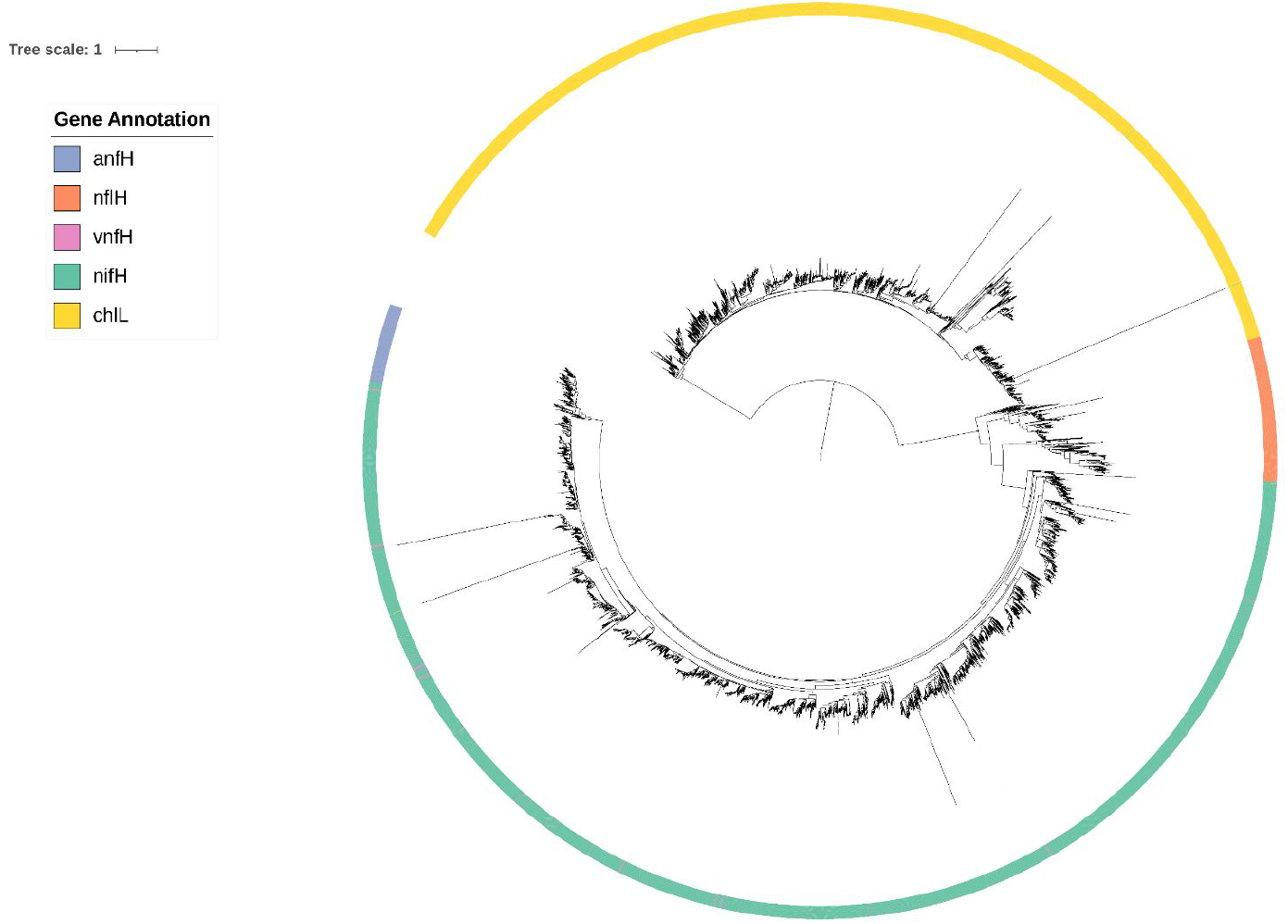
Gene tree of *nifH* and *nifH* homologs. The outer ring indicates final HMM annotations. Tree topology was used to estimate annotation consistency, with thirteen sequences assigned to a non-homogeneous clade (estimated error rate = 9.5 * 10^-4^).

**Figure S12:**
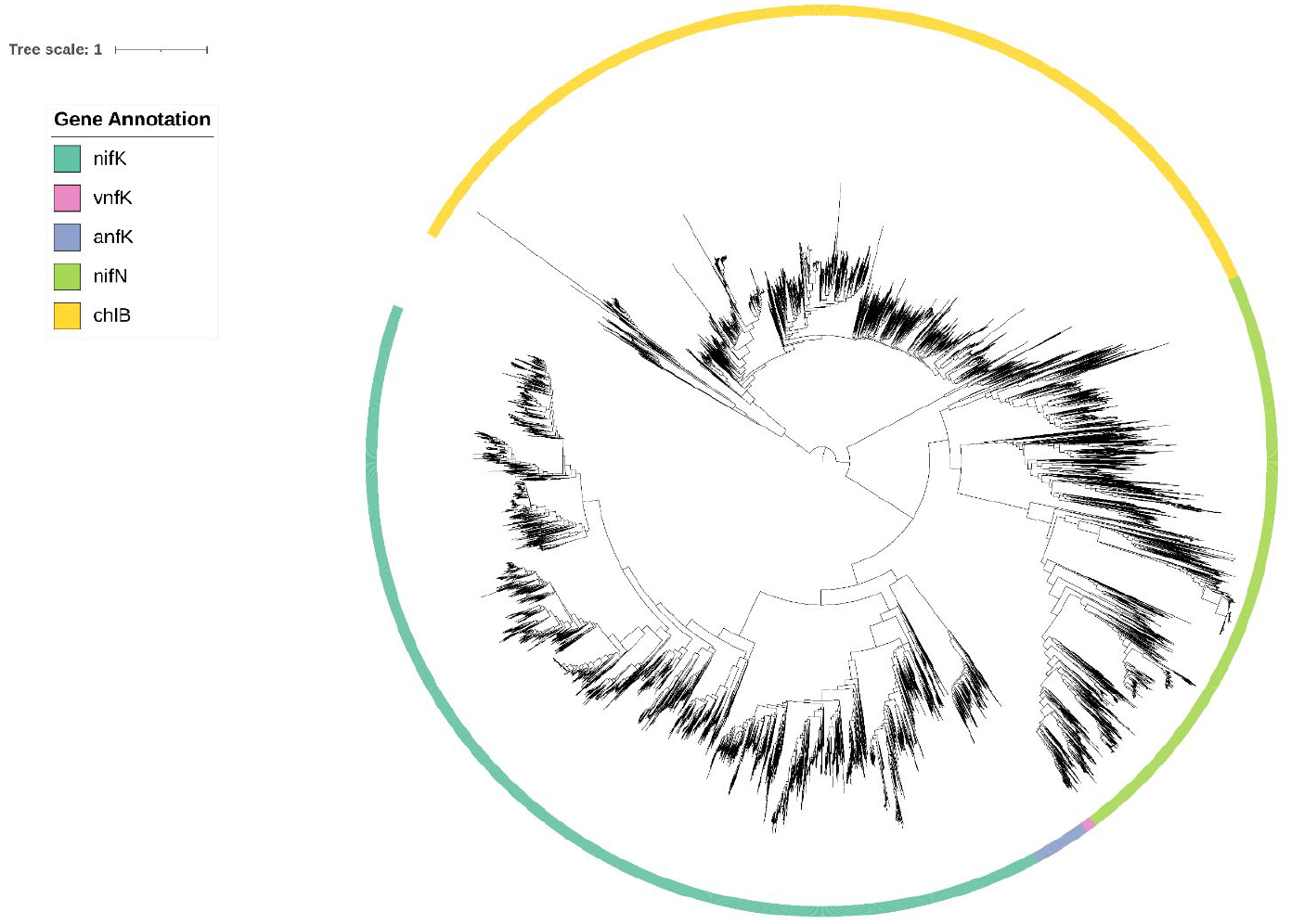
Gene tree of *nifK* and *nifK* homologs. The outer ring indicates final HMM annotations. Tree topology was used to estimate annotation consistency, with four sequences assigned to a non-homogeneous clade (estimated error rate = 1.8 * 10^-4^).

---

**Table S1:** Genes screened.

| Type | Genes |
| --- | --- |
| Molybdenum nitrogenase | *nifHDK* |
| Iron nitrogenase | *anfHDK* |
| Vanadium nitrogenase | *vnfHDK* |
| Nitrogenase biosynthetic genes | *nifEN* |
| Bacteriochlorophyll and chlorophyll biosynthesis (nitrogenase homolog) | *ChlLNB* |
| Nitrogenase like enzymes (nitrogenase homolog) | *nflHD* |

**Table S2:** Overview of all databases involved in the benchmarking analysis.

| DB | TYPE | ORIGIN (METHOD) / YEAR | SOURCE | AUTHORS |
| --- | --- | --- | --- | --- |
| **NFixPlanet** | Genome derived | Curated HMMs (fna) for *nif/vnf/anf HDK*, *nifEN*, *ChlLBN*, *nflHD*. Genomic context filters & tree based validation. | proGenomes3 SPIRE | This manuscript |
| **NFixDB** | Genome derived | Curated HMMs (faa) for *nif/vnf/anf HDK*, *ChlLBN*, *nflHD*. Genomic context filtering, 2024 | GTDB r214 | Bellanger et al. |
| **Turk-Kubo** | Amplicon | DADA2 pipeline for creating *nifH* ASV database (*nifH*1-4 primers only), data from 21 amplicon studies + 2 unpublished studies, 2024 | studies from NCBI SRA | Morando et al. |
| **Buckley** | Amplicon | Retrieval of *nifH* sequences via ARB + alignment, 2014 update | GenBank | Gaby and Buckley |
| **Zehr** | Amplicon | ARBitorator pipeline (BLASTx measure of similarity of gene to one of 15 representative *nifH* sequences), 2017 update | GenBank | Heller et al. |

**Table S3:** Fisher’s exact test for enrichment and depletion of *nifD* gene clusters across the ten phyla containing the greatest numbers of *nifD* clusters. Enrichment was assessed relative to the representation of each phylum in the combined proGenomes v3 and SPIRE genome databases. Odds ratios greater than one indicate enrichment, whereas odds ratios less than one indicate depletion.

| Phylum | Odds ratio | Enrichment p value | Depletion p value |
| --- | --- | --- | --- |
| Pseudomonadota | 2.07672397 | <2.2e-16 | 1 |
| Desulfobacterota | 30.1642514 | <2.2e-16 | 1 |
| Bacillota_A | 0.23328674 | 1 | <2.2e-16 |
| Bacteroidota | 0.33080319 | 1 | <2.2e-16 |
| Cyanobacteriota | 10.4447253 | <2.2e-16 | 1 |
| Verrucomicrobiota | 3.88401817 | <2.2e-16 | 1 |
| Campylobacterota | 1.62045663 | 7.73E-10 | 1 |
| Halobacteriota | 18.9047161 | <2.2e-16 | 1 |
| Acidobacteriota | 5.72214144 | <2.2e-16 | 1 |
| Nitrospirota | 26.4737749 | <2.2e-16 | 1 |

